# Multiplexed Single-cell Metabolic Profiles Organize the Spectrum of Cytotoxic Human T Cells

**DOI:** 10.1101/2020.01.17.909796

**Authors:** Felix J. Hartmann, Dunja Mrdjen, Erin McCaffrey, David R. Glass, Noah F. Greenwald, Anusha Bharadwaj, Zumana Khair, Alex Baranski, Reema Baskar, Michael Angelo, Sean C. Bendall

**Affiliations:** Department of Pathology, School of Medicine, Stanford University, Palo Alto, CA, 94305, USA; Immunology Graduate Program, Stanford University, Stanford, CA, 94305, USA

**Keywords:** Metabolism, Single-cell, Immunometabolism, Tumor Immunology, Cytotoxic T cells, Mass Cytometry, CyTOF, Multiplexed Ion Beam Imaging, MIBI-TOF

## Abstract

Cellular metabolism regulates immune cell activation, differentiation and effector functions to the extent that its perturbation can augment immune responses. However, the analytical technologies available to study cellular metabolism lack single-cell resolution, obscuring metabolic heterogeneity and its connection to immune phenotype and function. To that end, we utilized high-dimensional, antibody-based technologies to simultaneously quantify the single-cell metabolic regulome in combination with phenotypic identity. Mass cytometry (CyTOF)-based application of this approach to early human T cell activation enabled the comprehensive reconstruction of the coordinated metabolic remodeling of naïve CD8^+^ T cells and aligned with conventional bulk assays for glycolysis and oxidative phosphorylation. Extending this analysis to a variety of tissue-resident immune cells revealed tissue-restricted metabolic states of human cytotoxic T cells, including metabolically repressed subsets that expressed CD39 and PD1 and that were enriched in colorectal carcinoma versus healthy adjacent tissue. Finally, combining this approach with multiplexed ion beam imaging by time-of-flight (MIBI-TOF) demonstrated the existence of spatially enriched metabolic neighborhoods, independent of cell identity and additionally revealed exclusion of metabolically repressed cytotoxic T cell states from the tumor-immune boundary in human colorectal carcinoma. Overall, we provide an approach that permits the robust approximation of metabolic states in individual cells along with multimodal analysis of cell identity and functional characteristics that can be applied to human clinical samples to study cellular metabolism how it may be perturbed to affect immunological outcomes.

## Introduction

Immune cells dynamically execute highly context-dependent functions. During an immune reaction, cells rapidly respond to environmental triggers and migrate into affected tissues, expand exponentially and secrete large amounts of effector molecules to recruit and instruct other immune cells or kill invading pathogens. All of these diverse capacities are enabled and coordinated by dynamic changes in cellular metabolism^1–3^. Besides the generation of energy, metabolic intermediates provide critical building blocks for biosynthesis pathways as well as epigenetic and post-translational modifications, thus regulating gene expression and the generation of effector molecules^4,5^. Given these context-dependent roles of cellular metabolism, selective pharmacological targeting of individual pathways can influence specific aspects of immune cell behavior, for example direct the balance between effector and regulatory functionality^6,7^. Such therapeutic targeting has been shown to improve antitumor responses^8–10^, ameliorate autoimmune diseases^11,12^ and is a promising option for many other diseases^13^.

Central to many of these diseases, T cells constitute a heterogeneous population of specialized subsets with dedicated effector functions that are subject to metabolic influence^14,15^. Naïve T cells are characterized as metabolically quiescent, satisfying their minimal energy needs mostly through oxidative phosphorylation (OXPHOS). However, upon T cell receptor (TCR) engagement and appropriate co-stimulatory signaling, T cells drastically remodel their metabolism by increasing their engagement of aerobic glycolysis and amino acid metabolism^15^. Furthermore, differentiation and functional diversification of activated T cells are dependent on the engagement of specific metabolic programs, for example fatty acid metabolism which is important for the establishment of long-term memory T cell development^16^ and which has been shown to impact the balance between inflammatory effector T cells and suppressive regulatory T cells^17^. Many additional aspects of T cell biology, including T cell dysfunction and exhaustion^18^, are subject to such metabolic influence, together demonstrating the intimate connection between cellular metabolism and function which provides a unique opportunity to direct immune responses.

Approximation of the cellular metabolic state has been mostly based on quantification of metabolites and intermediates of specific metabolic pathways. Typically in bulk assays, mass spectrometry (MS) can be used to quantify metabolite abundances^19,20^ which can be further extended by introducing and tracing isotopically enriched metabolites, thus providing a method to track these compounds through a network of metabolic pathways^21^. Alternatively, an approach termed extracellular flux analysis allows the measurement of oxygen consumption and acidification of the extracellular milieu as proxies for OXPHOS and glycolytic activity, respectively^22,23^. Together, these technologies have yielded invaluable insight into cellular metabolism and they continue to provide the basis for many studies in the field of immunometabolism.

Still, significant challenges and open questions related to metabolic heterogeneity and its relationship with cell identity remain. Firstly, while several metabolic features have been shown to influence and direct T cell differentiation^24^, a more comprehensive and multimodal understanding of the timing and coordination within and between various metabolic pathways as well as the interplay with other cellular processes would allow to better direct T cell differentiation for various therapeutic uses. Furthermore, given the recently highlighted significant metabolic differences between physiologically activated cells and *in vitro* models^25^, there is a need to validate and more directly analyze metabolic states directly *ex vivo*. Especially analysis of limited samples from human clinical material could determine tissue-specific metabolic states as well as their potential modulation in human diseases, particularly cancer^26,27^. Moreover, multimodal analysis of metabolic immune cell states within their physiological microenvironment^3,28^ or in metabolically challenging contexts such as cancer could reveal novel therapeutic targets and guide clinical decisions^29,30^. An ideal technological solution to approach these questions would bridge the gap between highly multiplexed, single-cell phenotyping platforms and the bulk determination of metabolic state, thus enabling the study of cellular metabolism directly from *ex vivo* human clinical samples with sparse material while determining important metabolic and functional relationships^14^.

To address this need, we have developed a novel approach, termed single-cell metabolic profiling (scMEP), that enables quantification of single-cell metabolic states by capturing the composition of the metabolic regulome using antibody-based proteomic platforms such as mass cytometry^31^ and multiplexed ion beam imaging by time-of-flight (MIBI-TOF)^32^. These technologies make use of heavy metal-conjugated antibodies that are quantified by TOF MS, thus allowing highly multiplexed, single-cell and imaging assays^33^. First, we assessed over 110 antibody clones against metabolite transporters, metabolic enzymes, regulatory modifications (e.g. protein phosphorylation), signaling molecules and transcription factors across eight metabolic axes and on a variety of sample formats and tissue types. Utilizing these antibodies in multiplexed mass cytometry assays showed that heterogeneous populations such as human peripheral blood can be metabolically analyzed in a highly robust and repeatable manner and that cell identity is reflected in lineage-specific metabolic profiles. Using *in vitro*-activated human naïve and memory T cells, we benchmarked the scMEP approach against conventional extracellular flux analysis, demonstrating close agreement of metabolic target expression with glycolytic and respiratory activity. Multimodal analysis of a broad range of metabolic pathways and other cellular features during the metabolic switch of human naïve CD8^+^ T cells upon TCR engagement uncovered the coordinated nature of metabolic remodeling and its close connection to cell cycle progression, cell division and biogenesis. Furthermore, we investigated the tissue-specificity of metabolic profiles of human cytotoxic T cell subsets isolated from clinical samples, including colorectal carcinoma and healthy adjacent colon. This analysis revealed the metabolic heterogeneity of physiologically activated CD8^+^ T cell subsets, including subsets expressing the T cell exhaustion-associated molecules CD39 and PD1 that have recently been shown to predict therapeutic success^34^. Finally, we adopted the scMEP platform for the multiplexed imaging of human tissue samples by MIBI-TOF which revealed the spatial organization of metabolic T cell states in relation to the tumor-immune boundary as well as exclusion of clinically relevant CD8^+^ T cell subsets from this important microenvironment.

Overall, scMEP enables the study of cellular metabolic states in combination with phenotypic identity as well as effector functions. We expect this to deepen our understanding of cellular metabolism in homeostatic and dysfunctional settings, across heterogeneous cell populations and *in situ*, together providing a new lens through which to understand and impact human disease.

## Results

### Targeted quantification of the metabolic regulome discriminates human immune populations

Much like the epigenetic regulome governs transcription by controlling gene accessibility, the balance of cellular metabolites, and thus the metabolic state, is influenced by the molecular machinery that regulates these pathways. As such, we sought to quantify the abundance of metabolite transporters, rate-limiting metabolic enzymes and their regulatory modifications (e.g. phosphorylation), modifiers of mitochondrial dynamics^35^, as well as transcription factors and signaling molecules that drive specific metabolic programs^36^, here collectively referred to as the cellular metabolic regulome. Using a targeted, antibody-based approach to study a broad range of regulatory determinants enabled multimodal, single-cell analysis of the metabolic state in conjunction with other cell features (i.e. cell phenotype, cycling, signaling) that could be adopted to a multitude of high-dimensional probe-based technologies such as mass cytometry and MIBI-TOF (Fig. 1a).

**Fig. 1:**
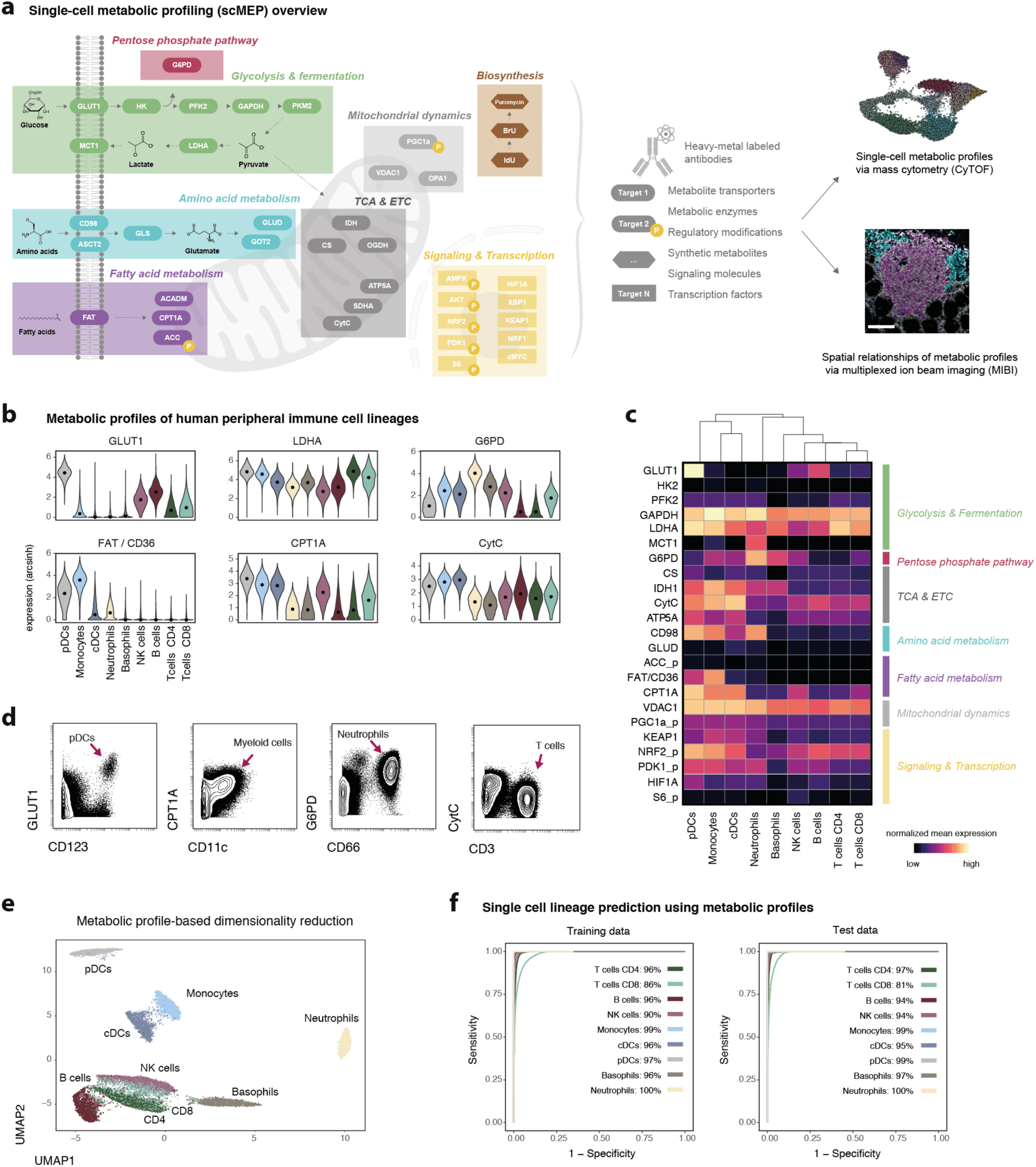
Single-cell metabolic profiles organize the human immune system. **a**, Conceptual overview of the employed scMEP approach. Important determinants and modifiers of metabolic activity were identified and respective probes (mostly, but not exclusively antibodies) were conjugated to heavy-metal isotopes for their use in mass cytometry (CyTOF) and MIBI-TOF. Various combinations of metabolic probes and antibodies against other cellular features of interest were employed across the different experiments in this study. For a full account of all probes tested in this study see Supplementary Table 1. Scale bar = 100 μm. **b**, Whole blood of healthy individuals (N = 5, for donor characteristics see Supplementary Table 2) was fixed and stained with an antibody panel of 23 metabolic and 22 immunological antibodies. Cell populations were identified through FlowSOM clustering and annotated into the major immune cell lineages. Shown are examples of (arcsinh transformed) expression values across identified peripheral immune cell lineages. Black dots represent population medians. **c**, Normalized (99.9^th^ percentile) mean expression of all assessed metabolic targets across immune cell lineages. **d**, Examples of metabolic target expression across immune cell lineages. Shown are live, single, CD45^+^ cells of one representative individual. **e**, Cells from all five donors were subsampled for equal representation of all immune cell lineages and all donors. Only metabolic profiles (23 targets) were used as input data to the UMAP-based dimensionality reduction. Cells are colored by their lineage identity determined as in b. **f**, L1 regularized linear regression (using only metabolic profiles) was trained on a subset of donor (N = 3) and tested on a separate set of donors (N = 2). Stated numbers report balanced accuracy for the indicated population.

To accomplish this, we first assessed assay-specific performance of over 110 commercially available antibodies (see Supplementary Table 1) following in-house heavy metal-conjugation. Tested antibodies targeted a wide range of metabolic pathways, including glycolysis and fermentation, amino acid metabolism, fatty acid metabolism as well as components of the tricarboxylic acid (TCA) cycle and electron transfer chain (ETC). The performance of individual heavy-metal conjugated antibodies was validated through mass cytometry, immunohistochemistry (IHC) and MIBI-TOF (Supplementary Fig. 1a,b), based on biological controls (cell lineage-specific expression patterns^37^ and induction upon activation^38,39^) but also inter-assay and inter-platform reproducibility and subcellular localization of targets. Following this screening, we selected a subset of metabolic antibodies (N = 41, Fig. 1a) that were used in varying combinations and in conjunction with antibodies against additional cellular characteristics for the experiments in this study.

Given the importance of metabolic networks to immune cell function, we first asked whether functional specialization within the human immune system might be reflected in lineage-specific metabolic profiles. We obtained whole blood from five healthy individuals (Supplementary Table 2) and, employing the outlined approach, simultaneously analyzed their single-cell metabolic profiles in combination with a range of lineage markers (Fig. 1b-d). Immune cell lineages were identified through FlowSOM clustering^40^ using their high-dimensional lineage marker expression patterns (CD45, CD3, CD4, CD8, CD45RA, CD66, CD14, CD19, CD20, HLA-DR, CD56, CD57, CD11c, CD123, FceRI, CD235ab; Supplementary Fig. 2a,b), thus enabling *in silico* subset selection and comparison of metabolic profiles of all identified human immune cell lineages without the need for prior isolation or enrichment which can influence metabolic states^41,42^.

We observed specific, lineage-associated metabolic profiles that were in agreement with previously established critical roles of these metabolic pathways in specialized immune cell functions (Fig. 1c,d). For example, plasmacytoid dendritic cells (pDCs) expressed the highest levels of the glucose transporter GLUT1 (also SLC2A1), high levels of the fatty acid translocase (FAT also CD36) and several other targets with glycolytic and fatty acid metabolism pathways. Both pathways have been shown to impact important aspects of pDC functionality, including hallmark interferon production upon TLR stimulation^43^. Further, neutrophils displayed high levels of glucose-6-phosphate dehydrogenase (G6PD), an integral part of the pentose phosphate pathway (PPP). G6PD deficiency has been shown to severely impact neutrophil functionality, resulting in increased susceptibility to infections^44^. In line with their metabolic quiescence in the absence of antigenic stimulation, lymphocytic immune cell populations expressed lower levels of many metabolic targets including those in the glycolytic pathway while expressing intermediate levels of proteins within the TCA and ETC, crucial for basal respiration.

Using these metabolic profiles, we calculated total metabolic distances (defined as Euclidean distance in the high-dimensional metabolic space) between human immune cell lineages, underlining the division of lymphocytic and myeloid cell lineages and their functional specialization (Supplementary Fig. 2c). Further, lineage-specific expression profiles were found to be reproducible across different donors as evident by metabolic profile-based principle component analysis (PCA) and hierarchical clustering (Supplementary Fig. 2d,e). Technical variability of our approach was assessed by performing an independent experiment with cells from the same five healthy donors, demonstrating high levels of inter-experiment agreement across metabolic targets (r^2^ = 0.99, Supplementary Fig. 2f,g). Furthermore, we analyzed longitudinal changes in metabolic profiles of peripheral blood immune cell lineages which were stored for 0-48 h at 4 °C before processing. Metabolic profiles were highly stable during this time-period, reflecting the suitability of the scMEP approach under common experimental settings (Supplementary Fig. 2h).

Given these robust and lineage-specific profiles, we hypothesized that scMEPs could be used to directly infer cell identity without the use of cell surface markers. We made use of the unsupervised UMAP dimensionality reduction approach^45^, using only metabolic profiles as input dimensions and disregarding typically employed surface/lineage markers (Fig. 1e). Metabolic profiles were largely able to separate immune cell lineages, with expected overlap between CD4^+^ and CD8^+^ T cells and to a lesser extent NK cells. Further, an L1 regularized classifier^46^ trained on cells from three of these donors (training data) and subsequently used to predict immune cell identity of cells from two separate donors (test data) based only on their metabolic profiles was able to correctly assign lineage identity to a large fraction of cells (95% average across populations in both, training and test data), with minor misclassifications between CD4^+^ and CD8^+^ T cells (Fig. 1f). Together, this demonstrates the suitability and robustness of the scMEP approach to study metabolic regulation and its relationship to single-cell specialization and it revealed remarkable cell type-specific metabolic diversification within the human immune system.

### Benchmarking single-cell metabolic profiles with metabolic flux during T cell activation

Assessing bulk metabolic enzyme abundances has been shown to provide important insight into the regulation of cellular metabolism in many scenarios^38,39,47^ and multiple studies have identified specific enzymes that determine metabolic pathway activity across many cell types^48,49^. To establish the relationship between antibody-based, single-cell metabolic profiles and pathway activity, we chose *in vitro* activation of human T cells as a model system to assess the predictive capacity of protein expression levels on glycolytic and respiratory activity (Fig. 2a). Human naïve and memory T cells (including CD4^+^ and CD8^+^) were isolated using negative magnetic selection and activated for 0 to 5 days with anti-CD3/anti-CD28 beads. Cells from all conditions were divided and analyzed by mass cytometry and extracellular flux analysis (Seahorse analysis system). Focusing on glycolytic and TCA/ETC components, mass cytometry-based scMEP analysis recapitulated several established hallmarks of metabolic remodeling upon TCR engagement^36,50,51^, including upregulation of surface expression of the glucose transporter GLUT1, as well as intracellular levels of several key glycolytic enzymes including hexokinase 2 (HK2), phosphofructokinase 2 (PFK2) and critical downstream enzymes such as lactate dehydrogenase A (LDHA) and the monocarboxylate transporter 1 (MCT1, Fig. 2b,c). Likewise, proteins within the TCA and the ETC such as citrate synthase (CS), oxoglutarate dehydrogenase (OGDH), the cytochrome complex (CytC) and ATP synthase (ATP5A) were upregulated upon TCR engagement (Fig. 2b,d).

**Fig. 2:**
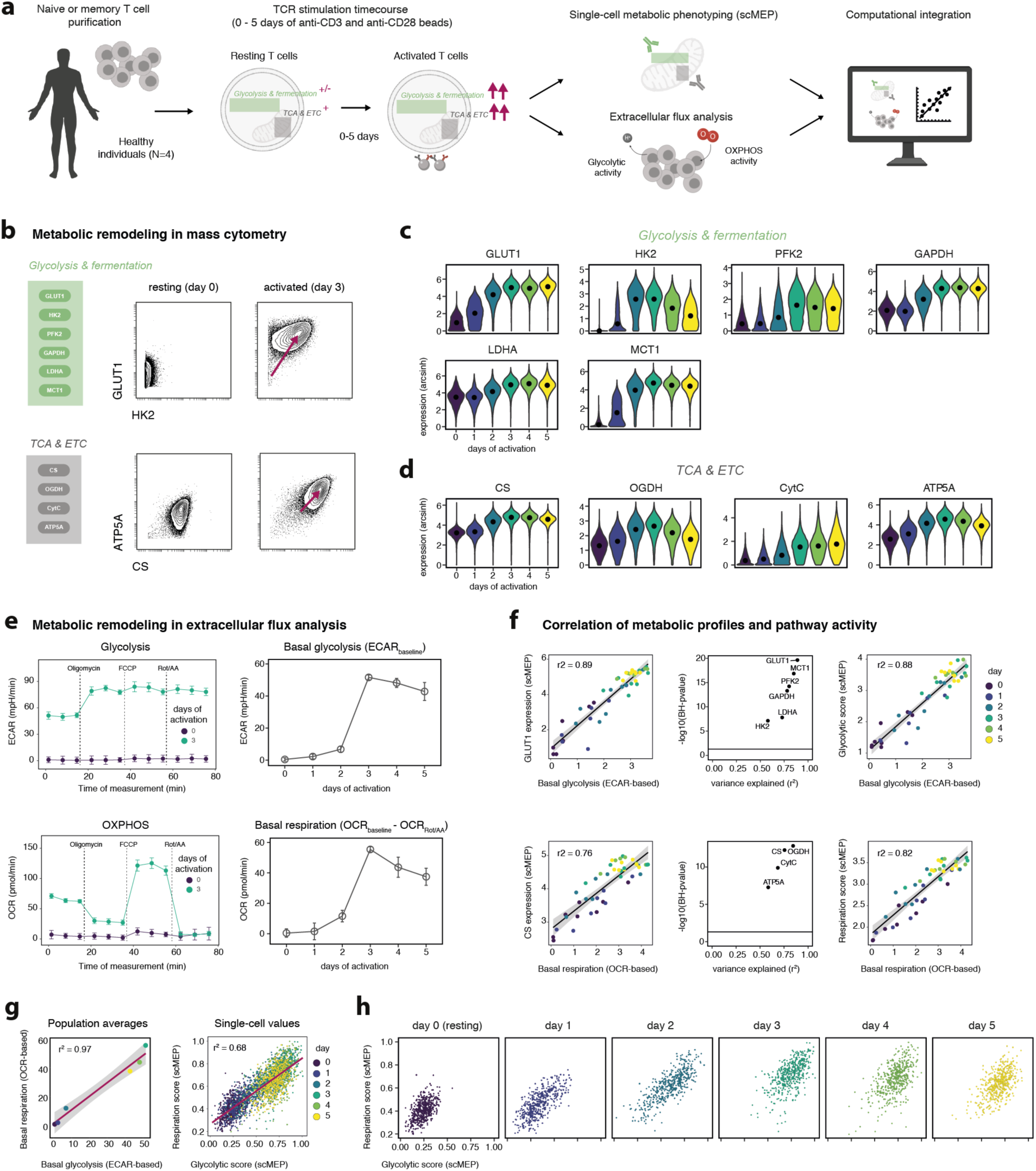
Single-cell metabolic profiles of T cell activation dynamics. **a**, PBMCs were isolated from healthy individuals (N = 4). Naïve or memory T cells (containing CD4^+^ and CD8^+^ cells) were purified by negative isolation with magnetic beads. Purified T cell populations (naïve or memory) were then activated using anti-CD3/anti-CD28 beads for 0-5 days. Cells from all conditions were divided into two samples: one sample was fixed and prepared for mass cytometry analysis while remaining cells were used for extracellular flux analysis in a Seahorse analyzer. Data from both modalities was then computationally integrated and compared. **b**, Examples of mass cytometry-quantified expression levels of important glycolytic (top) and TCA/ETC (bottom) components following no (resting, left) and 3 days (right) of anti-CD3/anti-CD28 activation on naïve T cells. Examples are gated on single, live, DNA^+^CD45^+^CD3^+^CD8^+^ cells, showing data of one representative experiment. **c**, Expression levels of important determinants of glycolysis/fermentation on naïve CD8^+^ T cells stimulated for 0-5 days. Black dots indicate population medians. **d**, Expression levels of ETC/TCA components on naïve CD8^+^ T cells as in c. **e**, Extracellular flux analysis of cells from the same experiments as in a-c. 100’000 naïve or memory T cells activated for 0-5 days were seeded into multiple wells. Extracellular acidification rate (ECAR; top) and oxygen consumption rate (OCR; bottom) for each measurement following injections of mitochondrial modifiers to determine basal pathway activity (right). FCCP = fluoro-carbonyl cynade phenylhydrazon, Rot = Rotenone, AA = antimycin A. Basal glycolysis = mean of the three baseline ECAR readings before injection. Basal respiration = mean baseline OCR – mean OCR post Rot/AA injection. Circles and error bars represent mean±s.d. **f**, Correlation between an exemplative (left panel) glycolytic and oxidative metabolic target and basal glycolysis (top row) and basal respiration (bottom row). Flux analysis values were arcsinh transformed (cofactor 5) for this analysis. Black lines and r^2^ values represent results of a linear regression model, with black shading representing the 95% CI. Log10 of (Benjamini-Hochberg) false discovery rate (FDR)-adjusted P-values (to control for multiple hypothesis testing) and r^2^ values from linear regression models (middle panel). Black line indicates a BH-corrected P-value of 0.05. Protein-based scMEP scores (right) represent the mean expression of all metabolic targets within a given pathway. Each dot represents the mean scMEP score of a T cell population (naïve or memory). **g**, Linear correlation of mean (left) and single-cell (right) OXPHOS and glycolytic scMEP scores of naïve CD8^+^ T cell populations, calculated as in f. **h**, scMEP scores as in g, visualized for each day of anti-CD3/anti-CD28 stimulation.

In order to benchmark these mass cytometry-based observations, we took the remaining T cells from the same experimental conditions and assessed their glycolytic and OXPHOS activity through extracellular flux analysis. As expected, levels of basal glycolysis and OXPHOS increased upon TCR engagement, peaking after 3-4 days of activation (Fig. 2e and Supplementary Fig. 3a). We used both bulk metabolic activities and single cell-derived mean metabolic target expression values for data integration and regression analysis (Fig. 2f). Glycolytic protein expression of all targets (GLUT1, HK2, PFK2, GAPDH, LDHA, MCT1) was robustly correlated with glycolytic flux values across several donors and independent experiments (linear regression mean r^2^ = 0.77) and a similarly strong correlation was observed between expression of TCA/ETC components (CS, OGDH, CytC, ATP5A) and OXPHOS activity (linear regression mean r^2^ = 0.72). We observed the highest r^2^ values for surface nutrient transporters (e.g. GLUT1, r^2^ = 0.89), potentially reflecting their previously demonstrated regulation through surface trafficking or internalization^52^. Downstream metabolic enzymes, whose enzymatic activity can additionally be modulated through post-translational modifications and feedback-inhibition^53^, nevertheless displayed substantial correlation values (e.g. PFK2 r^2^ = 0.80, HK2 r^2^ = 0.58).

Given this robust correlation of individual metabolic protein expression and pathway activity, we hypothesized that it would be possible to utilize scMEP-based high-dimensional co-expression patterns of multiple metabolic targets to derive *in silico* scores representing glycolytic and OXPHOS activity (Fig. 2f). We defined scores as the mean of (asinh-transformed and percentile-normalized) metabolic enzyme expression levels within a given pathway. Importantly, these scMEP-based glycolytic and OXHPOS scores strongly and robustly predicted respective metabolic activity across multiple donors, activation time points, and independent experiments as determined by linear regression (r^2^ = 0.88 and 0.82, respectively. Fig. 2f). Considering variability between different donors and experiments relevant to both mass cytometry and extracellular flux analysis, linear correlations of scMEP scores and pathway activity within each experiment displayed even stronger agreement (mean r^2^ = 0.92 and 0.86, respectively; Supplementary Fig. 3b).

Extracellular flux analysis suggested that upon TCR engagement, naïve T cell populations increase their glycolytic as well as their respiratory activity (r^2^ = 0.98; Fig. 2g). However, given the nature of bulk measurements, it remained unclear whether there is metabolic heterogeneity and specialization of a subset of cells towards a glycolytic or oxidative phenotype. Mass cytometry-based scMEP scores calculated for each cell independently indicated that, even on a single-cell level, cells simultaneously upregulate their glycolytic as well as oxidative machinery (r^2^ = 0.68; Fig. 2g), together supporting the notion that these cells simultaneously engage multiple metabolic pathways to support their wide-ranging bioenergetic demands^54^.

Furthermore, our single-cell analysis revealed previously obscured metabolic heterogeneity within each timepoint, with T cells activated for two days spanning almost the entire range of possible glycolytic and respiratory scMEP scores (Fig. 2h). Cells activated for three days displayed a more homogeneous upregulation of glycolytic and TCA/ETC enzymes, suggesting eventual convergence of metabolic remodeling and potentially indicating the presence of a series of metabolic and cellular checkpoints^55^. Together, this data validates the close relation of scMEP-based quantification of the cellular metabolic regulome with pathway activity assessed through a well-established orthogonal method and demonstrates the ability of scMEP to enable the discovery of metabolic heterogeneity at the single-cell level.

### Integrative modeling of T cell activation identifies checkpoints of metabolic switching

In order to more comprehensively study the metabolic remodeling of human T cells and its relation to cell activation, differentiation, and proliferation, we expanded our analysis to incorporate a broader set of metabolic pathways and other cellular features. In addition to glycolysis and OXPHOS (see Fig. 2), we included antibodies to analyze fatty acid and amino acid metabolism, both of which have been shown to impact T cell differentiation^36,56^, signaling molecules and transcription factors controlling metabolic networks, molecules regulating mitochondrial dynamics^35^, as well as anabolic metabolic activities spanning DNA, RNA and protein synthesis, monitored through the addition of the tagged substrates IdU, BrU and puromycin^57^. Importantly, we combined these metabolic measurements with analysis of critical cellular checkpoints, including immune cell activation, maturation and differentiation/exhaustion, cell cycle phase as well as cell division^58^. In total, we simultaneously analyzed 48 non-redundant, biological (non-technical) parameters across millions of single cells, allowing us to comprehensively study metabolic rewiring and its relationship to crucial cellular states (Fig. 3a).

**Fig. 3:**
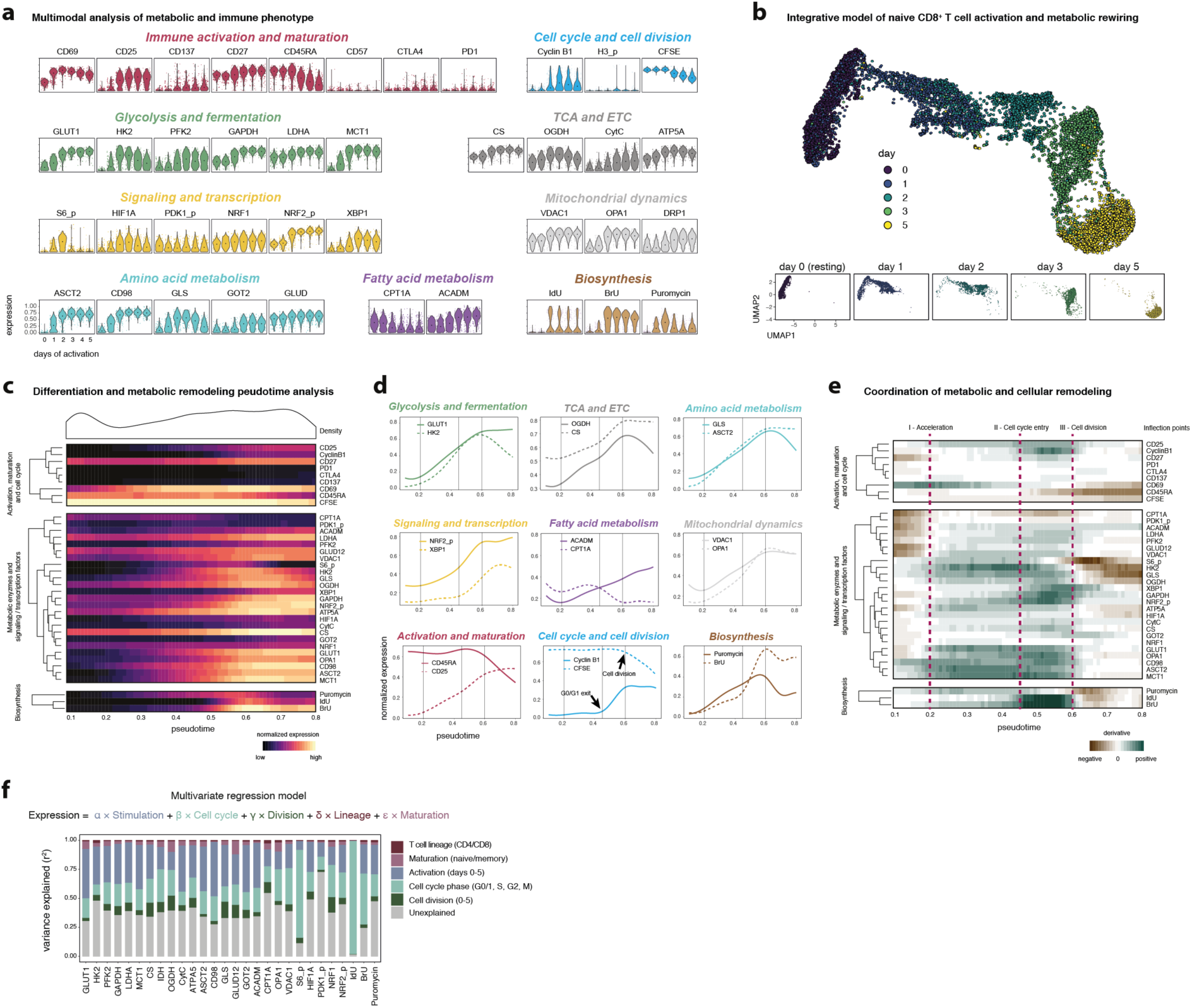
Integrative modeling of metabolic rewiring reveals determinants of human T cell activation. PBMCs were isolated from healthy individuals (N = 3). Naïve T cells (CD45RO^-^, containing CD4^+^ and CD8^+^ cells) were purified by negative isolation with magnetic beads. Purified T cell populations were then activated using anti-CD3/anti-CD28 beads for 0-5 days. **a**, Normalized (99.9^th^ percentile) expression of metabolic and phenotypic proteins by naïve CD8^+^ T cells across different days of activation. Shown are cells from one representative donor. Black dots indicate population medians. **b**, Cells were subsampled for equal representation of the indicated days of activation. Metabolic and other markers (with the exception of IdU and H3 phosphorylation) were used as input to UMAP dimensionality reduction and visualization. Cells are colored by their day of activation (top) and shown separately for each day of activation (bottom). **c**, Cells as in b were used as input to SCORPIUS trajectory inference using the same markers as in b. Data was grouped into 100 bins based on pseudotime. Heatmap depicts mean (scaled) expression levels of the indicated marker in the according pseudotime bin. Density (top) shows cell distribution along the pseudotime axis. **d**, Examples of continuous (smoothed) marker expression along pseudotime, calculated as in c and grouped by metabolic pathway. Vertical lines indicate important inflection points in the trajectory. Inflection points were chosen based on coordinated changes in the slope (see e) across the indicated metabolic markers and divide T cell metabolic remodeling into distinct stages. **e**, Slope (first derivative) of marker expression across pseudotime as in c. **f**, A multivariate regression model was used to quantify the variance explained (r^2^) by each of the indicated categories on metabolic marker expression. Shown are mean values from all four experiments.

As before, naïve human T cells were isolated using magnetic beads and activated for 0-5 days in vitro using anti-CD3/anti-CD28 beads. Indicating the initiation of metabolic remodeling, we observed early (within 24 h of activation) phosphorylation of signaling molecules (*e.g.* ribosomal protein S6) and induction of transcription factors (*e.g.* Hypoxia-inducible factor 1-alpha HIF1A). Upregulation of transcription factors and metabolite transporters was closely followed by upregulation of metabolic enzymes (*e.g.* sequential induction of S6 phosphorylation, HIF1A, GLUT1, HK2 and LDHA, Fig. 3a), together reminiscent of previously described temporal optimization patterns in which expression hierarchy matches enzyme order in metabolic pathways^59^. Indicating broader between-pathway coordination, other metabolic pathways were induced simultaneously, *e.g.* cells rapidly upregulated amino acid transporters ASCT2 and CD98/LAT1 as well as downstream glutaminase (GLS, Fig. 3a) which has been shown to be an important determinant for T cell differentiation^60^.

We next integrated the high-dimensional information from all metabolic and cellular features to calculate a two-dimensional UMAP projection of the metabolic and phenotypic progression of naïve CD8^+^ T cells upon TCR stimulation (Fig. 3b). This visualization indicated a continuous progression of immune activation and metabolic rewiring across multiple days with cells from early activation timepoints (24 and 48 h) spanning larger areas of the phenotypic and metabolic space compared to cells from later days (day 3 and 5). Indeed, activated naïve CD8^+^ T cells displayed the greatest metabolic heterogeneity (average metabolic cosine distance^58^) during early activation (days 0-2 average cosine distance = 0.07), converging during later days (days 3-5 average cosine distance = 0.038; Supplementary Fig. 4a). Consistent with previous reports^58^, these observations indicates that, like cell phenotype and cellular transcriptional profile, naïve CD8^+^ T cell metabolism is likely most plastic in the earliest phases of antigen experience.

To explore the temporal coordination of metabolic remodeling within and across metabolic pathways and in conjunction with cell phenotype, we made use of the SCORPIUS algorithm^61,62^ to infer a pseudotime axis representing progression of cellular differentiation and metabolic remodeling (Fig. 3c). Pseudotime correlated well with time of activation and was robust across different trajectory algorithms (SCORPIUS and Slingshot^63^) as well as independent biological replicates from healthy individuals (Supplementary Fig. 4b-e). We found metabolic protein expression within a given pathway to be highly coordinated (average within-pathway Spearman’s correlation r = 0.73, Supplementary Fig. 4f,g). Coordination was especially pronounced during early remodeling (pseudotime < 0.65; glycolysis r = 0.8, TCA/ETC r = 0.97, Amino acid r = 0.77). In addition to intra-pathway coordination, we found remarkable synchronization of metabolic protein expression trajectories across various metabolic and other cellular pathways (Fig. 3d,e). At later stages (pseudotime > 0.65), we observed several instances of divergent expression patterns (glycolysis r = -0.1, TCA/ETC r = 0.0, Amino acid r = -0.2, Supplementary Fig. 4f,g), potentially indicating redirection of metabolic intermediates into different metabolic pathways. For example, cells maintained high expression of the glutamine transporter ASCT2 but downregulated the glutamine-to-glutamate converting enzyme GLS, possibly increasing glutamine availability for nucleotide biosynthesis^64^.

Utilizing the change of metabolic protein expression over pseudotime (first derivative), we defined three inflection points during metabolic remodeling of naïve human CD8^+^ T cells, based on changes of the derivative of a broad range of metabolic markers (Fig. 3e). The first inflection point (pseudotime 0.2) was marked by a coordinated and accelerated upregulation of the cellular metabolic machinery in various pathways (e.g. concerted induction of GLUT1, ASCT2, OGDH, VDAC1 upregulation; Fig. 3e) leading up to the second inflection point (pseudotime 0.45) which was characterized by strong initiation of RNA synthesis (BrU incorporation), activation of the cellular stress response, including increased phosphorylation of the antioxidant transcription factor NRF2 [ref: ^65^] as well as upregulation of XBP1 which is part of the unfolded protein response and has been shown to control T cell function and mitochondrial activity^66^. Further, we observed reduced expression of carnitine palmitoyltransferase (CPT1A), a rate-limiting enzyme of fatty acid oxidation. Interestingly, this second inflection point coincides with cells exiting the G0/G1 cell cycle phase as evidenced by strong upregulation of DNA synthesis (IdU incorporation) and cyclin B1 expression (Fig. 3d,e). The third metabolic inflection point (pseudotime 0.6) was defined by stabilized or decreasing expression levels of various metabolic transporters and enzymes (e.g. GLUT1 and ASCT2) as well as peak translational activity (puromycin incorporation). Of note, this inflection point coincided with the first cell division (determined through reduction in CFSE signal). Interestingly, VDAC1 (also porin, located in the outer mitochondrial membrane) peaked at this point, suggesting an increase in mitochondrial mass prior to the first cell division, followed by cell division-dependent dilution^67^ and again underlining the crucial interplay between cell cycle progression, cell division and metabolic activity^68^.

Given this coordination of metabolic remodeling with cell activation, cell cycle progression and cell division, we sought to quantify their effects on metabolic protein expression using a multivariate regression model (Fig. 3f). We used time of activation (0-5 days), cell cycle phase (G0/G1, S, G2, M phase), number of cell divisions (0-5), T cell lineage (CD4/CD8) and maturation status (naïve/memory) as potential predictors of metabolic protein expression levels. Together, these determinants were able to account for 63% of the variance (mean r^2^) in metabolic marker expression. Activation was the single biggest determinant (r^2^ = 0.27), again demonstrating the extent of metabolic remodeling of T cells induced by TCR engagement. Activation was closely followed by cell cycle phase (r^2^ = 0.25) which, as expected had especially pronounced effects on biosynthesis pathways, but also on other metabolic pathways (e.g. cell cycle r^2^ = 0.22 for OGDH). Naïve and memory T cells have previously been shown to differentially engage metabolic pathways^15,69,70^ with memory T cells displaying increased basal glycolysis and respiration compared to naïve T cells (Supplementary Fig. 5a). Using scMEP, we confirmed memory CD8^+^ T cells to express higher levels of glycolytic (GLUT1 0.49 vs 0.14 median of memory vs. naïve, respectively) and TCA/ETC proteins (e.g. CS 0.66 vs. 0.54), increased levels of CPT1A (0.57 vs. 0.39) indicating increased fatty acid oxidation as well as elevated VDAC1 (0.52 vs. 0.38, Supplementary Fig. 5b) suggesting higher mitochondrial mass^71^. Integrating information from all metabolic markers further underlined differential metabolic profiles of resting memory and naïve T cells as shown by their distinct location on a two-dimensional UMAP projection (Supplementary Fig. 5c). Lastly, high-dimensional metabolic marker expression could be used to predict the cellular maturation state of resting CD8^+^ T cells (naïve vs memory, accuracy 97%; Supplementary Fig. 5d). Taken together, scMEP facilitated in-depth and multimodal analysis of early T cell activation which allowed us to reveal regulatory pathway coordination and to identify distinct phases of T cell metabolic remodeling as well as their relationship to transitions between cellular states.

### Tissue-specific metabolic profiles define human T cell subsets

In addition to cell-intrinsic factors, metabolic states are influenced by tissue-dependent factors such as nutrient availability^25^ and by microenvironmental modulations through malignantly transformed tumor cells^72^, further underlining the importance of studying metabolic states directly *ex vivo* from human biopsies and other clinical samples. To investigate these influences on cytotoxic T cells in the context of human cancer, we prepared single-cell suspensions from tissue resections of colorectal carcinoma patients, including sections from within the tumor (N = 6) but also adjacent healthy sections from the same patients (N = 6, see Supplementary Table 2). In addition, we included healthy donor PBMCs (N = 5) as well as lymph node biopsies (N = 3). All 20 samples were barcoded, combined into a single sample and stained with a panel of 18 phenotypic, as well as 27 metabolic antibodies (see Supplementary Table 1). We identified the main cell lineages and again confirmed their lineage-specific metabolic profiles (Supplementary Fig. 6a-e; see also Fig. 1).

Given their importance for anti-tumor immunity, we focused our downstream analysis on cytotoxic CD8^+^ T cells. We used FlowSOM and its meta-clustering functionality to group these cells based only on their high-dimensional co-expression patterns of metabolic markers, resulting in the definition of ten distinct metabolic scMEP states (Fig. 4a-c). Identified metabolic CD8^+^ T cell states included subsets characterized by low overall metabolic marker expression (scMEP 1&2) suggesting low metabolic activity, subsets with elevated expression of a broad range of targets (including CD98, GLUT1, PFK2, MCT1, CytC; scMEP 9&10), indicating increased metabolic demands, as well as subsets with characteristic expression patterns, e.g. increased ACADM/HIF1A (scMEP 4&7) or CD98 (scMEP 3&8; Fig. 4b,c) which might point to more specific, context-dependent metabolic adaptions.

**Fig. 4:**
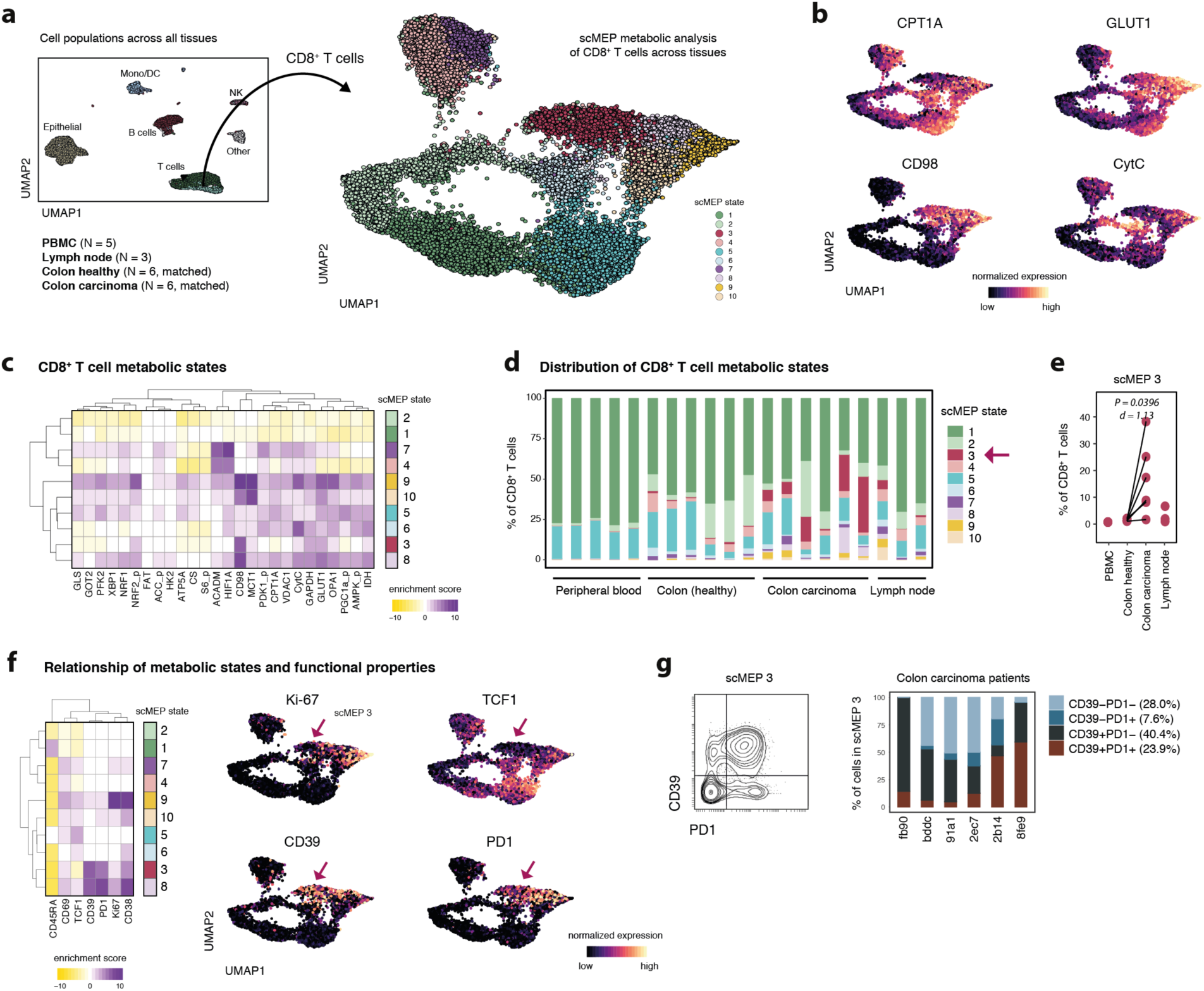
Cytotoxic T cell metabolic reflect tissue of residence. Healthy donor PBMC (N = 5), lymph node biopsies (N = 3) as well as single-cell suspensions from colorectal carcinoma (N = 6) and matched adjacent healthy sections (N = 6, see Supplementary Table 2) were barcoded, stained and acquired on a mass cytometer. **a**, Major cell lineages from all samples and tissues were identified through FlowSOM-based clustering. UMAP-dimensionality reduction was calculated using subsampled data from all lineages and all available markers. Cells are colored by their FlowSOM-based lineage definition (left). Next, total CD8^+^ T cells from all samples were selected and metabolic markers were used to define 10 scMEP states, based on FlowSOM clustering. UMAP-dimensionality reduction was calculated using subsampled data and only metabolic markers. Cells are colored by their scMEP state (right). **b**, UMAP visualization of CD8^+^ T cell scMEP states as in a colored by normalized (99.9^th^ percentile) expression of the indicated metabolic proteins. **c**, Marker enrichment modeling (MEM; ref^73^) was used to visualize enrichment (purple) or depletion (yellow) of metabolic target expression across CD8^+^ T cell scMEP states. **d**, Frequencies of scMEP state across individual samples. **e**, Statistical comparison of scMEP state frequencies (see also Supplementary Fig. 4f). P-values were calculated using a paired t-test between healthy and malignant sections from the same patient. Effect size is represented as Cohen’s d. **f**, MEM of immunological markers (not used for metabolic clustering) across scMEP states (left). UMAP visualization as in a,b with cells colored by their normalized expression value of the indicated immunological markers. **g**, Biaxial representation of cells from scMEP 3 pooled from all colorectal carcinoma samples (left). Frequencies of cells within scMEP 3, gated as PD1^+^ and CD39^+^ across all colorectal carcinoma samples (right).

Determining the tissue distribution of these phenotypes, we found that peripheral T cells primarily consisted of the metabolically low scMEP 1 (mean 76.6% of peripheral CD8^+^ T cells) with smaller numbers of scMEP 5 (19.7%) and negligible frequencies of all other metabolic states (all < 3%; Fig. 4d). Compared to peripheral blood, tissue infiltrating CD8^+^ T cells displayed higher metabolic heterogeneity with a more diverse range of phenotypes not found amongst peripheral blood cells, highlighting the importance of directly studying these cells using *ex vivo* assays. Further, this analysis allowed us to assess microenvironmental influences on T cell metabolism, specifically in malignant compared to healthy tissue. By comparing the relative frequency of each scMEP state between tissues (Supplementary Fig. 6f), we found a specific metabolic phenotype (scMEP 3) to be markedly enriched in cells isolated from colorectal carcinoma (mean 14.6% of total CD8^+^ T cells) versus cells from healthy adjacent sections (mean 0.8%) and all other tissues (PBMCs 0.1%, lymph nodes 2.4%; P = 0.04, Fig. 4e). Compared to other phenotypes, scMEP 3 was characterized by enriched expression of the amino acid transporter CD98 and lower levels of several metabolic enzymes (GLS, GOT2, PFK2, ATP5A, CS, S6_p, CPT1A, PGC1a_p), spanning a broad range of different pathways.

Since all CD8^+^ T cell scMEP states were defined using exclusively metabolic antibodies (all markers shown in the heatmap in Fig. 4c), we next investigated the relationship between metabolic profiles and immunological characteristics such as expression of activation/maturation markers, transcription factors and exhaustion-associated proteins (Fig. 4f). Importantly, scMEP metabolic states associated clearly with distinct immunological phenotypes. Metabolically low states (scMEP 1 & 5) displayed markers of resting cells (i.e. CD69, CD38 and Ki-67 low) with either a naïve (CD45RA and T cell factor 1 (TCF1) enriched, scMEP 1) or memory phenotype (CD45RA and TCF1 low, scMEP 5). Furthermore, the CD8^+^ T cell subpopulation with the highest expression of metabolic proteins (scMEP 9) showed indicators of recent activation and proliferation (i.e. CD69, CD38 and Ki-67 high), recapitulating the increased metabolic demands of actively cycling cells as also identified through our *in vitro* analysis of T cell differentiation (see Fig. 3).

Of note, the tumor-associated metabolic T cell state (scMEP 3) was significantly enriched in cells expressing the exhaustion-associated molecules programmed death 1 (PD1) and CD39 (both not used for the initial definition of scMEP states). Besides scMEP 3, we also observed enrichment of CD39^+^ and/or PD1^+^ cells (termed CD39/PD1 cells) in scMEP 8. With the exception of increased CD98 and minor enrichments in GLUT1 and GAPDH, cells from scMEP 3 but not scMEP 8, presented with decreased metabolic target expression (i.e. reduced ATP5A, CS, CytC and PGC1a phosphorylation) and were thus termed meta^low^ CD39/PD1 cells. Interestingly, lower mitochondrial capacity as indicated here has been shown to be characteristic of T cell exhaustion^8,74^. Furthermore, these meta^low^ CD39/PD1 cells (scMEP 3) displayed downregulated levels of TCF1 which has been proposed to be a hallmark feature of terminal exhaustion^75^. In comparison, cells from scMEP 8 retained TCF1 expression and were characterized by higher levels of many metabolic targets (termed meta^high^ CD39/PD1 cells), which might be indicative of different functional capacities of these subsets. Lastly, while the meta^low^ subset of tumor-infiltrating CD8^+^ T cells (scMEP 3) was strongly enriched in cells expressing CD39 and/or PD1 (mean 72% either CD39^+^ or PD1^+^), it also included a fraction of cells negative for both markers (28%, Fig. 4g), together suggesting that integration of the metabolic state could be employed as an additional dimension to functionally define T cell capacities. In summary, these analyses demonstrate the unique capability of scMEP to identify metabolic states of low abundance cell populations directly from sparse clinical samples, revealing important relationships between cellular phenotype and metabolism in human disease.

### Cellular metabolism is related to spatial organization in human tissue compartments

In addition to lineage intrinsic factors, cell activation and tissue of residence, a cell’s metabolism is influenced by its location within a specific tissue microenvironment and its interactions with neighboring cells^3,28^, together critically shaping immune responses against cancer^26,27^. The antibody-based nature of scMEP enabled us to directly transfer this approach to the recently-developed multiplexed ion beam imaging (MIBI-TOF) platform^32^, thus allowing multimodal, high-dimensional, image-based analysis of the cellular metabolic regulome, phenotypic state, and importantly, cellular interactions and localizations within the tissue microenvironment. In MIBI-TOF, a tissue section (e.g. FFPE tissue) is stained with heavy metal-conjugated antibodies and subsequently rastered with a primary ion beam, resulting in the release of secondary ions (including heavy-metals from target-bound antibodies). For each pixel, secondary ions are quantified by a TOF-MS system, thus allowing the reconstruction of high-dimensional images^33^.

To approach this, we first assessed the performance of all metabolic antibodies in traditional immunohistochemistry (IHC) versus MIBI-TOF (Supplementary Figs. 7 and 8). Next, we obtained human clinical tissue samples (i.e. archival FFPE blocks), including sections from colorectal carcinoma patients (N = 4) and non-matched non-malignant control sections (N = 3, different to patients analyzed in Fig. 4, see Supplementary Table 2). Tissue sections were stained with a combination of lineage markers^76^ and a broad range of metabolic antibodies (Fig. 5a and Supplementary Table 1). In total, we acquired 58 fields of view (FOV; 400 μm by 400 μm, resolution ∼400 nm), each comprised of 36 antibody-dimensions, thus allowing us to determine cell lineage, subset, and activation status, as well as metabolic state (Fig. 5b). Following pre-processing, deepcell^77^ was used to identify single cells within these images. Segmented cells were subsequently clustered into the main cell lineages (FlowSOM using vimentin, SMA, CD45, CD3, CD4, CD8, CD14, CD31, CD11c, CD68, CK, E-cadherin; Fig. 5c and Supplementary Fig. 8b,c). Importantly, throughout this process, the spatial location of each cell is retained, and cellular phenotypes can be investigated in their original tissue context (Supplementary Fig. 8d).

**Fig. 5:**
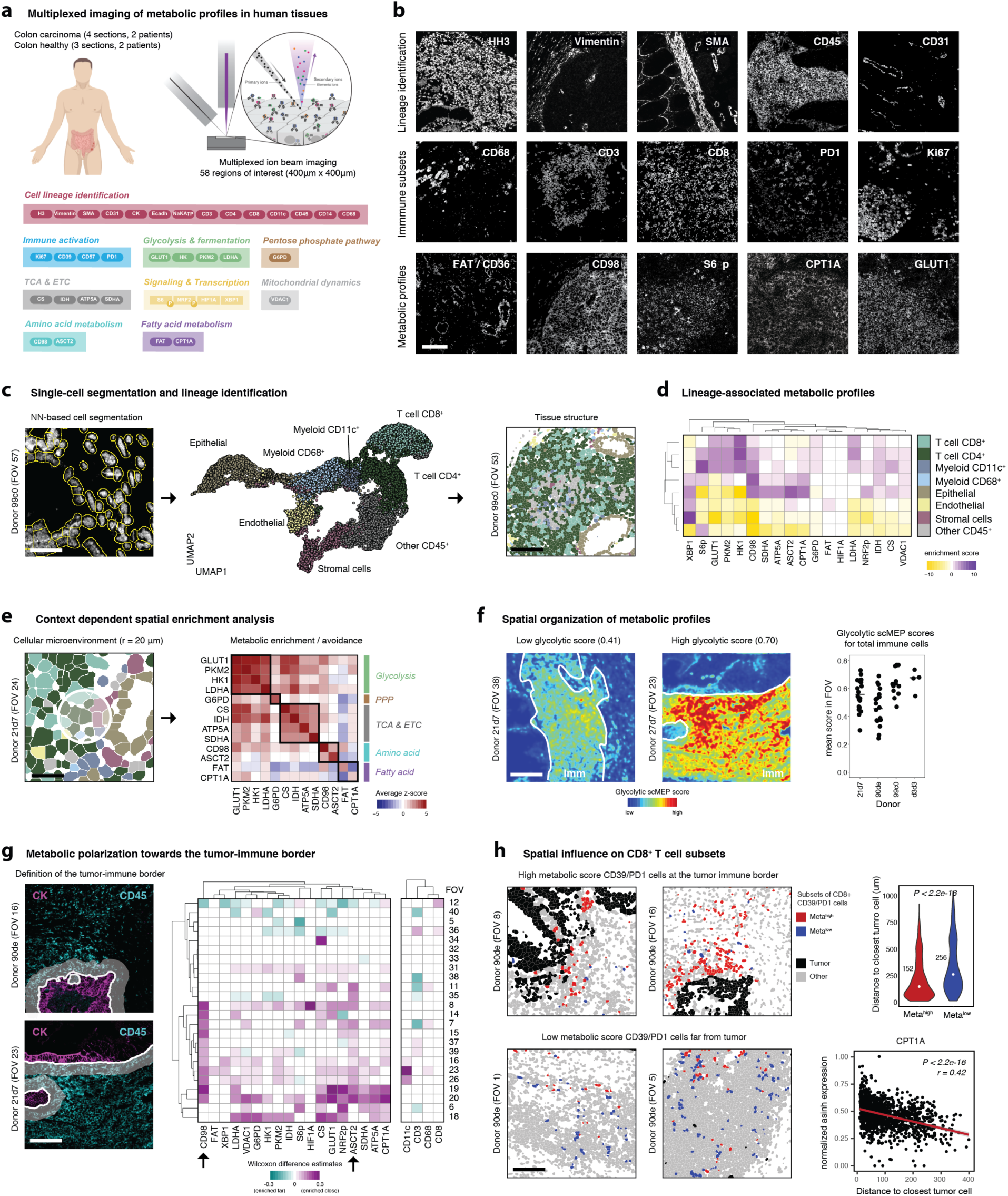
Imaging-based scMEP analysis reveals spatial influences on the organization of metabolic profiles in human colorectal carcinoma. **a**, FFPE colon-sections from colorectal carcinoma patients and healthy controls were stained with the indicated antibody panel (N = 36). A total of 58 fields of view (FOV), each 400 μm by 400 μm, were acquired by MIBI. **b**, Exemplary grayscale images of markers used for lineage identification (top), immune activation and subsets (middle) and their metabolic infrastructure (bottom). See also Supplementary Fig. 8. Scale bar = 100 μm. **c**, A pixel-based classifier was applied to automatically identify single-cells within these images (left, scale bar = 25 μm.). FlowSOM was used to identify the main cell lineages based on their lineage marker expression values. Single-cell data was projected onto two-dimensions using UMAP and colored by their cell lineage identity (middle). Clustered single-cell data can be mapped back onto the original segmented images to investigate spatial influences (right, scale bar = 100 μm). **d**, Metabolic profiles of cell lineages as identified in c are represented as MEM scores. **e**, Cellular microenvironments were defined as cells present within a 20 μm radius (based on cell centroids) of any given index cell. Colors indicate cell lineage as in d (left, scale bar = 25 μm). Within all such groups, spatial enrichments were calculated by comparing the distributions of metabolic marker expressions with a random subsampling of the same cell lineage composition. Enrichments (red) and avoidances (blue) are visualized as average z-scores across all FOVs. Black outlines indicate proteins within the same metabolic pathway (right). **f**, Spatial scMEP scores for a given metabolic pathway were calculated by averaging (and blurring) pixel-based expression values of all metabolic markers within a pathway. Areas of immune cell infiltration were outlined manually based on CD45 staining (left and middle, scale bar = 100 μm). Average glycolytic scMEP scores for all CD45^+^ cells within a FOV (right). **g**, The tumor-immune border was computationally defined using clustered and annotated single-cell data. Immune cells within a 20 μm radius of malignant epithelial cells were classified as located within the tumor-immune border (left, scale bar = 100 μm). Wilcoxon-rank sum test (adjusted for multiple hypothesis testing) were used to compare cells close to the border with cells further from the border in each FOV that contained cells of both categories. Heatmap shows Wilcoxon rank sum test-based estimates (representing the median of the difference between a samples from the two groups) of enrichment for enriched (magenta) and decreased (cyan) expression on immune cells within the border. Non-significant (BH-adjusted P-value > 0.05) estimates were colored white (right). **h**, CD8^+^ T cells expressing high levels of CD39 and/or PD1 were clustered into two subsets based on their metabolic target expression (see Supplementary Fig. 8). The two subsets (meta^high^ and meta^low^) were visualized in the original images (left, scale bar = 100 μm). Distance to closest malignant epithelial cell for CD39/PD1 cells stratified by metabolic phenotype. Numbers indicate median distance (top right). Linear regression of between normalized asinh expression of metabolic markers (e.g. CPT1A) and distance to closest tumor cell (bottom right). Red line and values indicate linear regression model.

To determine the metabolic influence of the local microenvironment and to reveal the spatial organization of metabolic states in colorectal carcinoma, we first used image-derived single-cell data to analyze cellular metabolic profiles (Fig. 5d). In agreement with our mass cytometry-based findings in peripheral blood and tissue-derived single-cell suspensions, distinct cell lineages displayed lineage-specific metabolic protein expression patterns with potentially activation-induced glycolytic expression in T cells and high metabolic protein levels in epithelial cells from colorectal carcinoma tissue.

Having established single-cell metabolic profiles consistent with our mass cytometry analysis, we next interrogated the spatial organization of metabolic profiles. For any given cell, we defined a cellular neighborhood consisting of cells found within a 20 μm distance to the center of the index cell (Fig. 5e) and then used context-dependent spatial enrichment (CDSE) analysis to determine if metabolic profiles are enriched in these neighborhoods^76^. CDSE compares the frequency of cells (within a certain radius) that express a given metabolic marker against random samplings of cells from the same lineage composition (for example same frequency of immune and epithelial cells), thus accounting for differential tissue structure, varying cell numbers and composition. We found that cells expressing high levels of specific metabolic targets were spatially enriched with cells displaying elevated levels of the same target. For example, GLUT1^high^ cells were highly enriched around other GLUT1^high^ cells, but also around cells with high levels other targets within the same metabolic pathway (e.g. GLUT1^high^ cells enriched around PKM2^high^ cells), thus suggesting the existence of environmental niches that enable or drive certain cellular metabolic behaviors irrespective of cell type and spanning healthy and colorectal carcinoma samples. We found such spatial enrichment for glycolysis, respiratory and amino acid pathways. However, FAT/CD36 and CPT1A, two targets involved in fatty acid metabolism were not spatially enriched. FAT/CD36 was specifically expressed on endothelial cells, potentially indicating their role in facilitating tissue uptake but not direct downstream oxidation of fatty acids^78^.

In analogy to single-cell mass cytometry scMEP scores (see Fig. 2), we calculated spatial scMEP scores from multiplexed images by averaging (and blurring) all images of markers within a given metabolic pathway, thus allowing visualization of their relative abundance related to spatial distributions (Fig. 5f and Supplementary Fig. 8e). Imaging several regions within a single tissue section, we found that average scMEP scores of total immune cells not only varied across donors but also within individual patients themselves (patient 21d7: 0.56±0.11, patient 90de: 0.46±0.13, mean±s.d.). At the same time, within a given immune cell infiltrate, we found indications of locally enriched metabolic scores (Fig. 5f, middle), altogether reinforcing the local, microenvironment-driven influence on metabolic polarization.

We hypothesized that malignant epithelial cells, known to be in metabolic competition with the immune system^49^, could constitute one such polarizing factor, locally influencing immune cell metabolism. To investigate this aspect, we identified immune cells close to the tumor-immune border (presence of epithelial cells within a 20 μm radius) and compared their metabolic state to immune cells located further away from the boundary^76^ (Fig. 5g). Here, a large fraction of FOVs (17 of 24 FOVs containing a tumor-immune boundary) displayed metabolic polarization towards the tumor region, dominated by increased expression of the amino acid transporters CD98 and ASCT2 (Fig. 5g, black arrows). Both of these transporters have been shown to be of prognostic value in various human cancers^79–81^ and it will be of great interest to see whether relation to tissue features such as the tumor border or multidimensional co-expression with other metabolic proteins will further improve their diagnostic power. Importantly, the observed metabolic polarization towards the tumor-immune border was not explained by variations in border immune cell lineage markers which were not significantly enriched across sampled regions (Fig. 5g, right panel of heatmap).

Finally, as suspension-based analysis of T cell metabolic diversity in colorectal carcinoma (see Fig. 4) had indicated, CD98 can be upregulated across various CD8^+^ T cell subsets, including two populations of CD39 and/or PD1 expressing cells (termed CD39/PD1) that differed in their metabolic profiles (meta^high^ and meta^low^). FlowSOM-based clustering of CD8^+^ CD39/PD1 cells within the imaging dataset using only their metabolic protein expression patterns again indicated the presence of two metabolically divergent subsets (meta^high^ and meta^low^, Fig. 5h and Supplementary Fig. 8g,h), akin to previously defined subsets. Investigating their spatial distribution, we found that CD39/PD1 meta^low^ cells were significantly further away from the tumor-border than their meta^high^ counterpart (meta^low^ 256 μm, meta^high^ 152 μm, *P* = 2.2e-16; Fig. 5h). In addition, we found a direct correlation (r = 0.42) between the distance to closest tumor cell and decreased expression of metabolic markers (most pronounced for CPT1A) in CD39/PD1 cells (Fig. 5h). Together, this analysis indicates that only CD39/PD1 cells that were distal and unengaged with the tumor appeared metabolically suppressed as opposed to the more metabolically active cells at the tumor-immune interface. These observations potentially reconcile the ambivalent nature of CD39 and PD1 surface expression, which is associated with exhaustion and dysfunction but at the same time T cell activation^82–84^. In summary, these spatial analyses revealed specific exclusion of metabolic immune cell subsets from the tumor-immune boundary, demonstrating the influence of tissue architecture on metabolic state that goes beyond what can be observed using conventional deep phenotyping of cell identity alone. Incorporating this new lens of single-cell metabolism into translational research promises better control of cellular alterations and dysfunction in human disease.

## Discussion

Biological tissues possess great functional and compositional heterogeneity, thus necessitating the use of single-cell platforms for their in-depth study^85,86^. Technological advances are pushing the frontier of such single-cell analyses on multiple cellular levels, including the genome^87,88^, transcriptome^89^ as well as aspects of the epigenome^90^ and proteome^31^. Here, we presented a novel approach termed single-cell metabolic profiling (scMEP) to investigate the cellular metabolic state that utilizes multiplexed, antibody-based assays to analyze metabolism-determining cellular features (Fig. 1). Focusing on the metabolic regulome allowed us to define single-cell metabolic states directly from limited, *ex vivo* human material. For example, we performed robust metabolic analyses of fewer than 1000 (median n=842) human tumor-infiltrating CD8^+^ T cells per sample (Fig. 4). Recent studies have highlighted significant metabolic differences between such physiologically activated cells and *in vitro* models^25^. Importantly, the scMEP approach is applicable to fixed cells and FFPE tissues, offering the opportunity to analyze metabolic states from existing clinical cohorts and thus enabling the identification of features associated with clinical outcome or therapeutic success.

Instead of individual metabolites, this scMEP approach quantifies the abundance of metabolic regulators such as metabolite transporters, rate-limiting metabolic enzymes as well as the activity of signaling pathways and transcription factors. In addition, we determined the biosynthesis rates of cellular macromolecules (RNA, DNA, protein) by quantifying the incorporation of BrU, IdU and puromycin, respectively^57^. Many of these factors have been previously shown to either directly drive or correlate with metabolic flux^36,39,48^ and we showed that scMEP metabolic profiles correlate robustly with metabolic activity as determined by extracellular flux analysis (Fig. 2). Nevertheless, scenarios might exist in which external factors (e.g. synthetic inhibitors) drive a divergence between the cellular need to activate metabolic programs and momentary pathway activity. Such inherently interesting exceptions would however be easily identified by subsequent validation of scMEP-based analyses with bulk MS or extracellular flux analysis.

Mass cytometry and MIBI-TOF both allow the quantification of >40 simultaneous features, allowing us to simultaneously analyze multiple major metabolic pathways. To do so, we validated a large number of commercially available antibodies (Supplementary Table 1) which provides resource for the implementation and potential future adjustments of scMEP, for example incorporation of additional metabolic pathways or comprehensive analysis of all components within a more restricted set of pathways. In addition, a large fraction (∼70%) of the tested metabolic antibodies are reactive with mouse and/or rat epitopes, thus facilitating straightforward transfer of this approach to analyze cells obtained from animal models. Of note, all antibodies were validated post cysteine-based heavy-metal conjugation which can potentially affect antibody-binding affinity. Presented antibody performances are therefore assay-specific and might reflect the impact of the employed conjugation chemistry.

Given the antibody-based nature of scMEP, this approach can be transferred to other high-dimensional probe-based platforms and could (with some limitations) also be employed in e.g. flow cytometry^33^. Furthermore, analysis of metabolic aspects using single-cell RNA sequencing in combination with novel analytical approaches could offer additional insights^91^. Especially once challenges related to RNA stability following cell fixation and permeabilization are resolved, the use of antibody-sequencing hybrid technologies^92,93^ would present an exciting platform to implement the scMEP approach, combining the unbiased nature of RNA sequencing with the large dynamic range of protein expression and the ability to assess post-transcriptional and post-translational regulation offered by antibody-based technologies^94^. In addition, metabolic profiling could be further extended by combining single-cell analysis of metabolic infrastructure and cellular phenotype with determination of epigenetic features^90^. Chromatin modification profiling has recently been demonstrated by mass cytometry^95^ and could be integrated with scMEP to further elucidate the reciprocal interplay of metabolism and epigenetic remodeling and it relation to human disease.

We demonstrated how employing scMEP can drive the discovery of biologically and relevant findings directly in clinical isolates. For example, comprehensive reconstruction of metabolic remodeling of T cells upon TCR engagement as demonstrated above (Fig. 3) could serve as a framework to design metabolic interventions in a phase-specific manner to direct *in vitro* differentiation of chimeric antigen receptor (CAR) T cells^96^ or cells used in adoptive cell transfer (ACT) therapy^97,98^. Furthermore, applying this approach to human clinical material revealed the presence of tissue-specific metabolic T cell subsets (Fig. 4). Here, we identified two metabolically diverging cytotoxic T cell subsets characterized by elevated expression of CD39 and PD1 and expanded in human colorectal carcinoma. CD39 and PD1 expression on immune cells has been shown to be clinically relevant in multiple tumor types^34,99^. Subsequent imaging analysis using the MIBI-TOF (Fig. 5) revealed that the metabolically-repressed CD39/PD1 cells were excluded from the tumor-immune boundary, a complex multicellular structure known to regulate immune function^76^. Importantly, while surface expression of CD39 and PD1 indicate T cell exhaustion/dysfunction, they can be expressed more broadly, driving the recent identification and integration of molecular regulators such as TOX^100–104^, NR4A^105,106^ and TCF1^75,107^ as more definitive indicators of immune cell dysfunction. Taken together, the here identified association of metabolic phenotype with CD39/PD1 expression and TCF1 downregulation, as well as the tumor-specific expansion of this metabolic subset in combination with its exclusion from the tumor-immune boundary suggest that incorporation of metabolic profiling to identify functionally diverse T cell states could further improve clinical stratification, e.g. to better predict response to immunotherapy.

In summary, we here presented a novel and robust approach to study single-cell metabolic states using antibody-based multiplex technologies. The application of scMEP should enable a better understanding of human immune cell biology and benefit the identification of disease-associated metabolic alterations that could serve as potential biomarkers and therapeutic targets for a variety of human diseases.

## Acknowledgements

We thank Leeat Keren and Jan van den Bossche for insightful discussions and invaluable feedback as well as the Nakamura lab at the Gladstone Institutes for access to their Seahorse XF analyzer. Further, we thank Albert Tsai for advice and help with clinical samples. F.J.H is supported by the EMBO organization (EMBO Long-Term Fellowship ALTF 1141-2017), the Novartis Foundation for medical-biological Research (16C148) and the Swiss National Science Foundation (SNF Early Postdoc Mobility P2ZHP3-171741). S.C.B. is supported by the NIH 1DP2OD022550-01, 1R01AG056287–01, 1R01AG057915-01 and 1U24CA224309-01.

## Author Contributions

Conceptualization, F.J.H., S.C.B.

Methodology, F.J.H., D.M., S.C.B.

Validation, F.J.H., D.M., A.B., Z.K.

Formal Analysis, F.J.H., D.R.G., E.M., N.G., A.B., D.M., R.B.

Investigation, F.J.H., S.C.B.

Resources, F.J.H., M.A., S.C.B.

Writing – Original Draft, F.J.H.

Writing – Review & Editing, F.J.H., S.C.B.

Visualization, F.J.H.

Supervision, F.J.H., S.C.B.

Project Administration, S.C.B.

Funding Acquisition, F.J.H., M.A., S.C.B.

## Methods

### Human samples

De-identified peripheral blood samples from healthy human subjects (see Supplementary Table 2) were obtained and experimental procedures were carried out in accordance with the guidelines of the Stanford Institutional Review Board (IRB). Written informed consent was obtained from all subjects. Fresh whole human blood in heparin collection tubes or leukoreduction system chamber contents (Terumo BCT) were obtained via the Stanford Blood Center. PBMCs were isolated via Ficoll (GE Healthcare) density gradient centrifugation.

FFPE tissue samples of colorectal carcinoma patients and healthy controls were obtained from the tissue repository of the Stanford Department of Pathology. Colorectal carcinoma and healthy adjacent tissue samples for mass cytometry (see Supplementary Table 2) were collected fresh after resection and transported for processing on ice in cell culture medium (CCM: RPMI-1640 (life technologies), 10% FBS, 1x L-glutamine, 1x penicillin/streptomycin (Thermo Fisher)). Samples were minced and processed using the MACS tumor dissociation kit (Miltenyi Biotec) as recommended. All viable single-cell suspensions were frozen in FBS supplemented with 10% DMSO and stored in liquid nitrogen.

### In vitro cell activation

Cryopreserved PBMC samples where thawed into 10 ml of cold CCM supplemented with 0.025 U/ml benzonase (Sigma) and washed once (250 g, 4 °C). Pan T cells were enriched through negative selection using magnetic beads (Pan T cell Isolation Kit, Miltenyi Biotec) as suggested by the supplier. Isolated T cells (including CD4^+^ and CD8^+^) were CFSE labeled by incubating them with 80 μm CSFE (Thermo Fisher) in CCM for 5 min at RT as described previously^58^. Labeled cells were quenched with warm CCM and washed three times by centrifuging for 5 min at 250 g. After washing, samples were divided for naïve and memory T cell isolation. Naïve T cells were enriched by depleting CD45RO expressing T cells using magnetic beads and memory T cells were isolated by depleting CD45RA expressing T cells. Using this approach, all cells used in in subsequent assays were negatively isolated. Cells were counted using an automated cell counting system and distributed into a 24-well plate in CCM at 1×10^6^ cells/well. For naïve T cells, 5 ng/ml of IL-2 was added to the culture and memory T cells were supplemented with 5 ng/ml IL-7 and IL-15 (all Peprotech). Cells were activated in a reverse time-course so that their total time in culture was identical and so that all cells would finish their indicated activation period at the same day to enable extracellular flux analysis. To do so, anti-CD3/anti-CD28 beads (Dynabeads, Thermo Fisher) were added in a 1:1 cell-to-bead ratio on the respective day and cells were incubated at 37 °C, 5% CO2 for up to 5 days. At the end of the activation period, cells from the same condition (same wells) were divided up and entered into the mass cytometry and extracellular flux analysis workflows.

### Live/dead discrimination and cell fixation

Cryopreserved single-cell suspensions (viable PBMC, lymph node and tumor biopsy samples) where thawed into 10 ml of cold CCM supplemented with 0.025 U/ml benzonase (Sigma) and washed once (250 g, 4 °C). *In vitro* activated T cell cultures (not cyropreserved) were washed once in CCM and directly processed further. For live / dead cell discrimination, monoisotopic cisplatin-194 (Fluidigm) was pre-conditioned for 48 h at 37 °C, aliquoted and stored at -20 °C. Viability staining was performed by resuspending cells in 1 ml of low-barium PBS and adding cisplatin-194 to a final concentration of 500 nM, followed by incubation for 5 min at RT and washing with cell staining medium (CSM: low-barium PBS with 0.5 % BSA and 0.02 % sodium azide (all Sigma)). Cells were fixed with 1.6% PFA in PBS for 10 min at RT and washed twice with CSM. Fixed cells were either entered directly into the staining workflow or cryopreserved by resuspending them in CSM supplemented with 10% DMSO and storing them at -80 °C.

### Heavy-metal conjugation of antibodies

Antibodies were conjugated to heavy-metal ions with MaxPar (Fluidigm) or MIBItag (Ionpath) reagents using an optimized conjugation protocol^108^. In short, antibodies were reduced with 4 mM TCEP (Thermo Fisher) for 30 min at 37 °C and washed two times. For conjugations using MaxPar reagents, metal chelation was performed by adding metal solutions (final 0.05 M) to chelating polymers and incubating for 40 min at RT. Metal-loaded polymers were washed twice with using a 3 kDa MWCO microfilter (Millipore) by centrifuging for 30 min, 12’000 g at RT. For conjugations using MIBItag reagents, pre-loaded polymers were obtained, and no loading reactions needed to be performed. For both approaches, antibody buffer exchange was performed by washing purified antibody through a 50 kDa MWCO microfilter (Millipore) and centrifuging for 10 min, 12’000 g at RT. Partially-reduced antibodies and metal-loaded polymers were incubated together for 90 min at 37 °C. Conjugated antibodies were washed four times and collected by two centrifugations (2 min, 1’000 g, RT) into an inverted column in a fresh 1.6 ml collection tube. Protein content was assessed by NanoDrop (Thermo Fisher) measurement, antibody stabilization buffer (Candor Bioscience) was added to a final volume of at least 50 v/v % and antibodies were stored at 4 °C.

### Palladium barcoding and staining with heavy-metal conjugated antibodies

To eliminate technical variability during staining or acquisition, individual samples within one experiment were palladium-barcoded as described previously^109^ and combined into a composite sample before further processing and staining. Cell-surface antibody master-mix in CSM was filtered through a pre-wetted 0.1 μm spin-column (Millipore) to remove antibody aggregates and added to the samples. After incubation for 30 min at RT, cells were washed once with CSM. To enable intracellular staining, cells were permeabilized by incubating with ice-cold MeOH for 10 min on ice and washed to times with CSM to remove any residual MeOH. Intracellular antibody master-mix in CSM was added to the samples and incubated for 1 h at RT. Cells were washed once with CSM and resuspended in intercalation solution (1.6% PFA in PBS and 0.5 μM rhodium-intercalator (Fluidigm)) for 20 min at RT or overnight at 4 °C.

Before acquisition, samples were washed once in CSM and twice in ddH_2_O and filtered through a cell strainer (Falcon). Cells were then resuspended at 1 x 10^6^ cells/mL in ddH_2_O supplemented with 1x EQ four element calibration beads (Fluidigm) and acquired on a CyTOF2 mass cytometer (Fluidigm).

### Antibody validation workflow

To validate metabolic antibodies as broadly as possible, a range of different cell types, tissues and technologies were used in combination. Mass cytometry-based antibody validation was performed on a range of cell lines, immune populations found in whole blood and T cells with or without TCR activation. First, various leukemic, embryonic and carcinoma cell lines were cultured in standard conditions^110^, fixed, palladium-barcoded and subsequently stained with heavy-metal conjugated antibodies as described (see above). Whole blood was processed as before (see above) and stained with a combination of metabolic antibodies and cell lineage markers (CD45, CD3, CD4, CD8, CD45RA, CD66, CD14, CD19, CD20, HLA-DR, CD56, CD57, CD11c, CD123, FceRI, CD235ab [ref: ^111^]) to identify the major immune cell types through manual gating. Human T cells were either rested or activated with anti-CD3/anti-CD38-beads for 72 h (see above), fixed and palladium-barcoded before staining with metabolic antibodies. Metabolic antibodies were initially used at a concentration of 2 μg/ml. For all populations, median arsinh values were calculated and positive staining was defined as a median of at least 10 ion counts (asinh transformed value >1.5) of any subpopulation. Where available, cell-lineage specific expression and induction upon activation were compared to previously determined values for the given cell population^37–39^.

To validate antibodies on tissues, control tonsil and liver FFPE tissues were stained with the indicated metal-conjugated antibodies as described below and their performance was validated through traditional IHC and MIBI-TOF. Detectable staining was determined through visual inspection of both IHC and grayscale MIBI-TOF images. For intra-assay quality control, IHC and MIBI-TOF images were visually compared and in addition, related to previously determined staining patterns^37^.

### Extracellular flux analysis

Extracellular flux analysis was performed by adopting previously outlined protocols^112^. In short, *in vitro* activated T cells were spun onto a Cell-Tak (Thermo Fisher) coated XF96 cell culture microplate (Agilent) with a density of 100’000 or 150’000 cells/well and rested in Seahorse XF RMPI 1640 medium supplemented with 2 mM L-glutamine, 2 mM sodium pyruvate and 25 mM glucose (all Agilent) for 1 h in a non-CO2 incubator at 37 °C. ECAR and OCR were measured using a Seahorse XF96 extracellular flux analyzer (Agilent). Oligomycin (1 μM), fluoro-carbonyl cyanide phenylhydrazone (FCCP; 1.5 μM) and rotenone (0.5 μM) together with antimycin A (0.5 μM) were sequentially injected to establish baseline parameters. Raw data was imported into the R environment^113^ in order to calculate basal glycolysis respiration rates as described previously^22^. Data was normalized by cell number. For linear regression between extracellular flux analysis values and mass cytometry values, both were arcsinh transformed with a cofactor of 5.

### Mass cytometry data preprocessing

Raw mass cytometry data was first bead-normalized to remove acquisition-related influences on marker expression using the premessa R package. Next, barcoded cells were assigned back to their initial samples using their unique palladium barcode combination. Normalized data was then uploaded onto cytobank.org^114^ or cellengine.com to identify single, live cells by manually gating on DNA (103Rh) and viability (194Pt) channels. Data was subsequently imported into the R environment, arcsinh transformed (cofactor 5) and normalized to the 99.9^th^ percentile of each respective channel before downstream analysis.

### Clustering and data visualization

Pre-processed single-cell (mass cytometry and MIBI-TOF) data was clustered using the FlowSOM R package^40^ and the indicated input channels. Resulting clusters were either manually annotated with the main cell lineages based on their lineage marker profiles or, if the underlying number and identity of clusters was unknown (e.g. for clustering on metabolic profiles) the metaclustering function of the FlowSOM package was used. Given their widespread expression across cell lineages, differences in metabolic target expression between different populations and clusters were visualized by marker enrichment modeling^73^. UMAP embeddings were calculated using the R uwot implementation with the following parameters: n_neighbors = 15, min_dist = 0.02.

### Calculation of metabolic scMEP scores

To calculate single-cell scMEP scores, expression values (debarcoded, bead-normalized, arcsinh transformed and percentile-normalized as described above) from all metabolic enzymes within a given pathway (glycolysis, respiration, amino acid metabolism and fatty acid metabolism) were summed and divided by the number of channels within the pathway.

To calculate image-based scMEP scores, pixel-based expression values from pre-processed data were blurred with a gaussian filter (sigma = 6) and arcsinh transformed. Next, pixel values from images within a given pathway were summed and finally percentile normalized to the 99^th^ percentile.

### Trajectory analysis of metabolic remodeling

Pre-processed data was randomly subsampled to represent all indicated days of activation equally. Pseudotime was calculated using the SCORPIUS^61^ and Slingshot^63^ algorithms, given their documented robustness across different datasets^62^, making use of the dynverse R implementation. All indicated channels were used as input dimensions to both algorithms and we did not define priors. Mitotic (M phase) cells and were excluded from this analysis given their drastically different metabolic profile. Strongly cell phase-dependent markers (IdU incorporation and H3 phosphorylation^115^) were not used as input dimensions for the trajectory calculation. Resulting pseudotime was scaled from 0-1. While resting cells (day 0) and cells from day 5 were included in the trajectory calculation to allow identification of starting and end points, we were mostly interested in early T cell activation (pseudotime 0.1-0.8), thus focusing downstream analysis on this period which additionally constitutes the most robust part of the trajectory as determined by comparison between SCORPIUS and Slingshot trajectories.

### Multivariate regression model

In order to assess the influence of different cellular and experimental conditions on metabolic profile expression, we designed a multivariate regression model using the relaimpo R implementation^116^. As input data, we used mean population expression values of all metabolic markers. Potential predictors were time of activation (0-5 days), cell cycle phase (G0/G1, S, G2, M phase, identified through manual gating^115^), number of cell divisions (0-5, identified through CFSE dilution), T cell lineage (CD4, CD8, identified through manual gating) and T cell maturation status (naïve, memory, separated through magnetic enrichment). Data from different donors was analyzed separately and displayed as averages across experiments.

### Analysis of scMEP repeatability and robustness

To determine the robustness of the scMEP approach, metabolic marker expression values of samples from the same healthy donors were stained and analyzed in two separate experiments and compared by linear regression using the lm() function. Hierarchical clustering using the R function hclust() was performed using the same input data. For the training phase of immune cell lineage prediction from metabolic profiles, 20’000 cells were randomly subsampled from three healthy donors to create an L1 regularized linear regression model as described before^95^ using the glmnet R package^46^. For the test phase, data was derived from two independent healthy donors not included in the training data. Prediction of T cell maturation status from metabolic profiles was performed similarly by randomly subsampling cells from both conditions into separate training and test data. Metabolic heterogeneity was defined as the cellular Euclidean distance to the average expression levels of the given population^58^. Only (pre-processed) metabolic profile expression values were used for this calculation.

### Staining for multiplexed ion beam imaging

Tissue sections (4 μm) were cut from colorectal carcinoma and control colon FFPE tissue blocks using a microtome and mounted on silanized gold-coated slides (IONpath) for MIBI-TOF analysis. Mounted tissue sections were incubated at 70°C for 20 min and deparaffinized with three washes of fresh xylene followed by rehydration with successive washes of ethanol 100% (2x), 95% (2x), 80% (1x), 70% (1x), and distilled water. Washes were performed using a Leica ST4020 Linear Stainer (Leica Bio-systems) programmed to three dips per wash for 30 s each. Rehydrated sections were immersed in epitope retrieval buffer (Target Retrieval Solution, pH 9, DAKO Agilent), incubated at 97 °C for 40 min and cooled down to 65 °C using Lab vision PT module (Thermofisher Scientific). Slides were washed with MIBI wash buffer (low-barium PBS IHC Tween buffer (Cell Marque) containing 0.1% (w/v) BSA (Thermofisher Scientific)). Slides were then placed into a Sequenza staining rack (Thermofisher Scientific) and sections were blocked for 1 h with BBDG blocking buffer (1X TBS IHC Wash Buffer with Tween 20 (Cell Marque) + 2% donkey serum, 0.1% cold fish skin gelatin (Sigma), 0.1% Triton X-100, and 0.05% sodium azide). Metal-conjugated antibody mix was prepared in 3% (v/v) donkey serum TBS IHC wash buffer and filtered using a centrifugal filter with a 0.1 mm PVDF membrane (Ultrafree-MC, Merck Millipore). Sections were stained with the antibody mix, incubating overnight at 4°C in the Sequenza staining rack. After overnight incubation, slides were washed twice with MIBI wash buffer and fixed for 5 min in 2% glutaraldehyde solution (Electron Microscopy Sciences) in low-barium PBS. Slides were then rinsed briefly in low-barium PBS and then dehydrated with successive washes of Tris 0.1 M (pH 8.5) (3x), distilled water (2x), and ethanol 70% (1x), 80%(1x), 95% (2x), 100% (2x). Slides were immediately dried in a vacuum chamber for at least 1 h prior to imaging.

### Immunohistochemistry

All MIBI-TOF antibodies were validated by DAB chromogenic IHC. The protocol for IHC closely followed the MIBI-TOF staining protocol, with minor changes. Before assembly of slides into the Sequenza staining rack and blocking, endogenous peroxidase activity was quenched by incubation in 3% H_2_O_2_ for 30 min and sections were washed with H_2_O on an orbital shaker for 5 min. Sections were stained with MIBI antibodies individually, and detected with ImmPRESS universal (Anti-Mouse/Anti-Rabbit) secondary antibody kit (Vector labs) and ImmPACT DAB Substrate kit (Vector Labs), according to the manufacturer’s guidelines.

### Multiplexed ion beam imaging acquisition

Quantitative imaging was performed using a custom designed MIBI-TOF mass spectrometer (IONpath), as previously described^32,76^, with an image size of 400 μm^2^ and 1024 x 1024 pixels. The entire cohort of 58 FOVs was acquired over a 24 h period of continuous imaging, yielding a total of 2088 images.

### Imaging data pre-processing and single-cell segmentation

Multiplexed imaging data was preprocessed as described before^76^. In short, for each pixel, MS spectra were converted into pixel counts by extracting a mass range from atomic mass unit amu-0.25 to amu±0. Background (due to absence of tissue or high gold signal) was removed, and noise was filtered out using a k-nearest-neighbor approach. To segment single cells from images, we trained a convolutional neural network^76,117^ using annotated training data from a variety of different cancer types. The network output was fed into the watershed algorithm to produce individual cells. This mask was used to extract per-cell counts for each marker in each image. Counts were normalized by cell size to account for different sampling of cells in the given plane. Normalized data was imported into the R environment and transformed using an inverse hyperbolic sine (arcsinh) cofactor of 0.05 (adjusted due to cell size normalization).

### Context-dependent spatial enrichment analysis

The context-dependent spatial enrichment (CDSE) approach^76^ was used to identify structured patterns of metabolic protein expression in the tissue. For each pair of metabolic markers, X and Y, the number of times cells positive for protein X was within a 50 pixel (∼20 um) radius of cells positive for protein Y was counted. Thresholds for positivity were customized to each marker individually. A null distribution was produced by performing 1000 bootstrap permutations where the locations of cells positive for protein Y were randomized. Randomizations retained the distribution of cells positive for protein Y across major lineage categories: immune, endothelial, epithelial, and fibroblast. A z-score was calculated comparing the number of true cooccurrences of cells positive for protein X and Y relative to the null distribution. For each pair of metabolic proteins X and Y the average z-score was calculated across malignant and control tissues separately.

### Visualization

Plots were created using the ggplot2 R package^118^. Schematic representations were created with biorender (https://biorender.io/). Figures were prepared in Illustrator (Adobe).

### Data availability

Mass cytometry: All data will be made available through flowrepository.org Multiplexed ion beam imaging: All data will be made available through ionpath.com

## References

1. Klein Geltink, R. I., Kyle, R. L. & Pearce, E. L. Unraveling the Complex Interplay Between T Cell Metabolism and Function. Annu. Rev. Immunol. 36, 461–488 (2018).

2. Olenchock, B. A., Rathmell, J. C. & Vander Heiden, M. G. Biochemical Underpinnings of Immune Cell Metabolic Phenotypes. Immunity 46, 703–713 (2017).

3. Buck, M. D., Sowell, R. T., Kaech, S. M. & Pearce, E. L. Metabolic Instruction of Immunity. Cell 169, 570–586 (2017).

4. Peng, M. et al. Aerobic glycolysis promotes T helper 1 cell differentiation through an epigenetic mechanism. Science 354, 481–484 (2016).

5. Chang, C.-H. et al. Posttranscriptional Control of T Cell Effector Function by Aerobic Glycolysis. Cell 153, 1239–1251 (2013).

6. Xu, T. et al. Metabolic control of TH17 and induced Treg cell balance by an epigenetic mechanism. Nature 548, 228–233 (2017).

7. Patel, C. H., Leone, R. D., Horton, M. R. & Powell, J. D. Targeting metabolism to regulate immune responses in autoimmunity and cancer. Nat. Rev. Drug Discov. 18, 669–688 (2019).

8. Scharping, N. E. et al. The Tumor Microenvironment Represses T Cell Mitochondrial Biogenesis to Drive Intratumoral T Cell Metabolic Insufficiency and Dysfunction. Immunity 45, 374–388 (2016).

9. Kishton, R. J., Sukumar, M. & Restifo, N. P. Metabolic Regulation of T Cell Longevity and Function in Tumor Immunotherapy. Cell Metab. 26, 94–109 (2017).

10. Leone, R. D. et al. Glutamine blockade induces divergent metabolic programs to overcome tumor immune evasion. Science 366, 1013–1021 (2019).

11. Angiari, S. et al. Pharmacological Activation of Pyruvate Kinase M2 Inhibits CD4+ T Cell Pathogenicity and Suppresses Autoimmunity. Cell Metab. 0, (2019).

12. Kornberg, M. D. et al. Dimethyl fumarate targets GAPDH and aerobic glycolysis to modulate immunity. Science 360, 449–453 (2018).

13. Lee, C.-F. et al. Preventing Allograft Rejection by Targeting Immune Metabolism. Cell Rep. 13, 760–770 (2015).

14. Pearce, E. L., Poffenberger, M. C., Chang, C.-H. & Jones, R. G. Fueling Immunity: Insights into Metabolism and Lymphocyte Function. Science 342, 1242454 (2013).

15. Buck, M. D., O’Sullivan, D. & Pearce, E. L. T cell metabolism drives immunity. J. Exp. Med. 212, 1345–1360 (2015).

16. van der Windt, G. J. W. et al. Mitochondrial Respiratory Capacity Is a Critical Regulator of CD8+ T Cell Memory Development. Immunity 36, 68–78 (2012).

17. O’Neill, L. A. J., Kishton, R. J. & Rathmell, J. A guide to immunometabolism for immunologists. Nat. Rev. Immunol. 16, 553–565 (2016).

18. MacIver, N. J., Michalek, R. D. & Rathmell, J. C. Metabolic Regulation of T Lymphocytes. Annu. Rev. Immunol. 31, 259–283 (2013).

19. Dettmer, K., Aronov, P. A. & Hammock, B. D. Mass spectrometry-based metabolomics. Mass Spectrom. Rev. 26, 51–78 (2007).

20. Gowda, G. A. N. & Djukovic, D. Overview of mass spectrometry-based metabolomics: opportunities and challenges. Methods Mol. Biol. 1198, 3–12 (2014).

21. Buescher, J. M. et al. A roadmap for interpreting 13C metabolite labeling patterns from cells. Curr. Opin. Biotechnol. 34, 189–201 (2015).

22. Mookerjee, S. A., Gerencser, A. A., Nicholls, D. G. & Brand, M. D. Quantifying intracellular rates of glycolytic and oxidative ATP production and consumption using extracellular flux measurements. J. Biol. Chem. 292, 7189–7207 (2017).

23. Nicholas, D. et al. Advances in the quantification of mitochondrial function in primary human immune cells through extracellular flux analysis. PLoS One 12, e0170975 (2017).

24. Verbist, K. C. et al. Metabolic maintenance of cell asymmetry following division in activated T lymphocytes. Nature 532, 389–393 (2016).

25. Ma, E. H. et al. Metabolic Profiling Using Stable Isotope Tracing Reveals Distinct Patterns of Glucose Utilization by Physiologically Activated CD8+ T Cells. Immunity 51, 856–870.e5 (2019).

26. O’Sullivan, D., Sanin, D. E., Pearce, E. J. & Pearce, E. L. Metabolic interventions in the immune response to cancer. Nat. Rev. Immunol. 19, 324–335 (2019).

27. Li, X. et al. Navigating metabolic pathways to enhance antitumour immunity and immunotherapy. Nat. Rev. Clin. Oncol. 16, 425–441 (2019).

28. Muir, A. & Vander Heiden, M. G. The nutrient environment affects therapy. Science 360, 962–963 (2018).

29. Kim, J. & DeBerardinis, R. J. Mechanisms and Implications of Metabolic Heterogeneity in Cancer. Cell Metab. 30, 434–446 (2019).

30. Le Bourgeois, T. et al. Targeting T Cell Metabolism for Improvement of Cancer Immunotherapy. Front. Oncol. 8, 237 (2018).

31. Bendall, S. C. et al. Single-cell mass cytometry of differential immune and drug responses across a human hematopoietic continuum. Science 332, 687–96 (2011).

32. Keren, L. et al. MIBI-TOF: A multiplexed imaging platform relates cellular phenotypes and tissue structure. Sci. Adv. 5, eaax5851 (2019).

33. Hartmann, F. J. & Bendall, S. C. Immune monitoring using mass cytometry and related high-dimensional imaging approaches. Nat. Rev. Rheumatol. (2019) doi:10.1038/s41584-019-0338-z.

34. Sade-Feldman, M. et al. Defining T Cell States Associated with Response to Checkpoint Immunotherapy in Melanoma. Cell 176, 404 (2019).

35. Buck, M. D. et al. Mitochondrial Dynamics Controls T Cell Fate through Metabolic Programming. Cell 166, 63–76 (2016).

36. Wang, R. et al. The Transcription Factor Myc Controls Metabolic Reprogramming upon T Lymphocyte Activation. Immunity 35, 871–882 (2011).

37. Uhlen, M. et al. Tissue-based map of the human proteome. Science 347, 1260419–1260419 (2015).

38. Howden, A. J. M. et al. Quantitative analysis of T cell proteomes and environmental sensors during T cell differentiation. Nat. Immunol. 20, (2019).

39. Geiger, R. et al. L-Arginine Modulates T Cell Metabolism and Enhances Survival and Anti-tumor Activity. Cell 167, 829–842.e13 (2016).

40. Van Gassen, S. et al. FlowSOM: Using self-organizing maps for visualization and interpretation of cytometry data. Cytometry. A 87, 636–45 (2015).

41. Binek, A. et al. Flow Cytometry Has a Significant Impact on the Cellular Metabolome. J. Proteome Res. 18, 169–181 (2019).

42. Llufrio, E. M., Wang, L., Naser, F. J. & Patti, G. J. Sorting cells alters their redox state and cellular metabolome. Redox Biol. 16, 381–387 (2018).

43. Saas, P., Varin, A., Perruche, S. & Ceroi, A. Recent insights into the implications of metabolism in plasmacytoid dendritic cell innate functions: Potential ways to control these functions. F1000Research 6, 456 (2017).

44. Gray, G. R. et al. Neutrophil dysfunction, chronic granulomatous disease, and non-spherocytic haemolytic anaemia caused by complete deficiency of glucose-6-phosphate dehydrogenase. Lancet 2, 530–4 (1973).

45. McInnes, L. & Healy, J. UMAP: Uniform Manifold Approximation and Projection for Dimension Reduction. arXiv (2018).

46. Friedman, J., Hastie, T. & Tibshirani, R. Regularization Paths for Generalized Linear Models via Coordinate Descent. J. Stat. Softw. 33, 1–22 (2010).

47. Rakus, D., Gizak, A., Deshmukh, A. & Wiśniewski, J. R. Absolute Quantitative Profiling of the Key Metabolic Pathways in Slow and Fast Skeletal Muscle. J. Proteome Res. 14, 1400–1411 (2015).

48. Tanner, L. B. et al. Four Key Steps Control Glycolytic Flux in Mammalian Cells. Cell Syst. 7, 49–62.e8 (2018).

49. Chang, C.-H. et al. Metabolic Competition in the Tumor Microenvironment Is a Driver of Cancer Progression. Cell 162, 1229–1241 (2015).

50. Finlay, D. K. et al. PDK1 regulation of mTOR and hypoxia-inducible factor 1 integrate metabolism and migration of CD8 + T cells. J. Exp. Med. 209, 2441–2453 (2012).

51. Frauwirth, K. A. et al. The CD28 Signaling Pathway Regulates Glucose Metabolism. Immunity 16, 769–777 (2002).

52. Wieman, H. L., Wofford, J. A. & Rathmell, J. C. Cytokine stimulation promotes glucose uptake via phosphatidylinositol-3 kinase/Akt regulation of Glut1 activity and trafficking. Mol. Biol. Cell 18, 1437–1446 (2007).

53. Wilson, J. E. Isozymes of mammalian hexokinase: structure, subcellular localization and metabolic function. J. Exp. Biol. 206, 2049 LP – 2057 (2003).

54. Slack, M., Wang, T. & Wang, R. T cell metabolic reprogramming and plasticity. Mol. Immunol. 68, 507–512 (2015).

55. Wang, R. & Green, D. R. Metabolic checkpoints in activated T cells. Nat. Immunol. 13, 907–915 (2012).

56. Sinclair, L. V et al. Control of amino-acid transport by antigen receptors coordinates the metabolic reprogramming essential for T cell differentiation. Nat. Immunol. 14, 500–508 (2013).

57. Kimmey, S. C., Borges, L., Baskar, R. & Bendall, S. C. Parallel analysis of tri-molecular biosynthesis with cell identity and function in single cells. Nat. Commun. 10, 1185 (2019).

58. Good, Z. et al. Proliferation tracing with single-cell mass cytometry optimizes generation of stem cell memory-like T cells. Nat. Biotechnol. 37, 259–266 (2019).

59. Zaslaver, A. et al. Just-in-time transcription program in metabolic pathways. Nat. Genet. 36, 486–491 (2004).

60. Johnson, M. O. et al. Distinct Regulation of Th17 and Th1 Cell Differentiation by Glutaminase-Dependent Metabolism. Cell 175, 1780–1795.e19 (2018).

61. Cannoodt, R. et al. SCORPIUS improves trajectory inference and identifies novel modules in dendritic cell development. bioRxiv 079509 (2016) doi:10.1101/079509.

62. Saelens, W., Cannoodt, R., Todorov, H. & Saeys, Y. A comparison of single-cell trajectory inference methods. Nat. Biotechnol. 37, 547–554 (2019).

63. Street, K. et al. Slingshot: cell lineage and pseudotime inference for single-cell transcriptomics. BMC Genomics 19, 477 (2018).

64. Altman, B. J., Stine, Z. E. & Dang, C. V. From Krebs to clinic: glutamine metabolism to cancer therapy. Nat. Rev. Cancer 16, 619–634 (2016).

65. Ma, Q. Role of Nrf2 in Oxidative Stress and Toxicity. Annu. Rev. Pharmacol. Toxicol. 53, 401–426 (2013).

66. Song, M. et al. IRE1α–XBP1 controls T cell function in ovarian cancer by regulating mitochondrial activity. Nature 562, 423–428 (2018).

67. Fischer, M. et al. Early effector maturation of naïve human CD8 ^+^ T cells requires mitochondrial biogenesis. Eur. J. Immunol. 48, 1632–1643 (2018).

68. Icard, P., Fournel, L., Wu, Z., Alifano, M. & Lincet, H. Interconnection between Metabolism and Cell Cycle in Cancer. Trends Biochem. Sci. 44, 490–501 (2019).

69. Gerriets, V. A. & Rathmell, J. C. Metabolic pathways in T cell fate and function. Trends Immunol. 33, 168–173 (2012).

70. van der Windt, G. J. W. & Pearce, E. L. Metabolic switching and fuel choice during T-cell differentiation and memory development. Immunol. Rev. 249, 27–42 (2012).

71. van der Windt, G. J. W. et al. CD8 memory T cells have a bioenergetic advantage that underlies their rapid recall ability. Proc. Natl. Acad. Sci. 110, 14336–14341 (2013).

72. Renner, K. et al. Metabolic Hallmarks of Tumor and Immune Cells in the Tumor Microenvironment. Front. Immunol. 8, 248 (2017).

73. Diggins, K. E., Greenplate, A. R., Leelatian, N., Wogsland, C. E. & Irish, J. M. Characterizing cell subsets using marker enrichment modeling. Nat. Methods 14, 275–278 (2017).

74. Bengsch, B. et al. Bioenergetic Insufficiencies Due to Metabolic Alterations Regulated by the Inhibitory Receptor PD-1 Are an Early Driver of CD8 + T Cell Exhaustion T Cell Exhaustion. Immunity 45, 358–373 (2016).

75. Blank, C. U. et al. Defining ‘T cell exhaustion’. Nat. Rev. Immunol. 19, 665–674 (2019).

76. Keren, L. et al. A Structured Tumor-Immune Microenvironment in Triple Negative Breast Cancer Revealed by Multiplexed Ion Beam Imaging. Cell 174, 1373–1387.e19 (2018).

77. Van Valen, D. A. et al. Deep Learning Automates the Quantitative Analysis of Individual Cells in Live-Cell Imaging Experiments. PLoS Comput. Biol. 12, e1005177 (2016).

78. Son, N.-H. et al. Endothelial cell CD36 optimizes tissue fatty acid uptake. J. Clin. Invest. 128, 4329–4342 (2018).

79. Kaira, K. et al. Prognostic significance of l-type amino acid transporter 1 (LAT1) and 4F2 heavy chain (CD98) expression in stage I pulmonary adenocarcinoma. Lung Cancer 66, 120–126 (2009).

80. Furuya, M., Horiguchi, J., Nakajima, H., Kanai, Y. & Oyama, T. Correlation of L-type amino acid transporter 1 and CD98 expression with triple negative breast cancer prognosis. Cancer Sci. 103, 382–389 (2012).

81. Shimizu, K. et al. ASC amino-acid transporter 2 (ASCT2) as a novel prognostic marker in non-small cell lung cancer. Br. J. Cancer 110, 2030–2039 (2014).

82. Canale, F. P. et al. CD39 Expression Defines Cell Exhaustion in Tumor-Infiltrating CD8+ T Cells. Cancer Res. 78, 115–128 (2018).

83. Raczkowski, F. et al. CD39 is upregulated during activation of mouse and human T cells and attenuates the immune response to Listeria monocytogenes. PLoS One 13, e0197151 (2018).

84. Simon, S. & Labarriere, N. PD-1 expression on tumor-specific T cells: Friend or foe for immunotherapy? Oncoimmunology 7, e1364828 (2018).

85. Heath, J. R., Ribas, A. & Mischel, P. S. Single-cell analysis tools for drug discovery and development. Nat. Rev. Drug Discov. 15, 204–216 (2016).

86. Wang, D. & Bodovitz, S. Single cell analysis: the new frontier in ‘omics’. Trends Biotechnol. 28, 281–290 (2010).

87. Kalisky, T. & Quake, S. R. Single-cell genomics. Nat. Methods 8, 311–314 (2011).

88. Linnarsson, S. & Teichmann, S. A. Single-cell genomics: coming of age. Genome Biol. 17, 97 (2016).

89. Papalexi, E. & Satija, R. Single-cell RNA sequencing to explore immune cell heterogeneity. Nat. Rev. Immunol. 18, 35–45 (2018).

90. Schwartzman, O. & Tanay, A. Single-cell epigenomics: techniques and emerging applications. Nat. Rev. Genet. 16, 716–726 (2015).

91. Xiao, Z., Dai, Z. & Locasale, J. W. Metabolic landscape of the tumor microenvironment at single cell resolution. Nat. Commun. 10, 3763 (2019).

92. Stoeckius, M. et al. Simultaneous epitope and transcriptome measurement in single cells. Nat. Methods 14, 865–868 (2017).

93. Shahi, P., Kim, S. C., Haliburton, J. R., Gartner, Z. J. & Abate, A. R. Abseq: Ultrahigh-throughput single cell protein profiling with droplet microfluidic barcoding. Sci. Rep. 7, 44447 (2017).

94. Mair, F. et al. A targeted multi-omic analysis approach measures protein expression and low abundance transcripts on the single cell level. bioRxiv 700534 (2019) doi:10.1101/700534.

95. Cheung, P. et al. Single-Cell Chromatin Modification Profiling Reveals Increased Epigenetic Variations with Aging. Cell 173, 1385–1397.e14 (2018).

96. Sadelain, M. Chimeric Antigen Receptors: A Paradigm Shift in Immunotherapy. Annu. Rev. Cancer Biol. 1, 447–466 (2017).

97. Rosenberg, S. A. & Restifo, N. P. Adoptive cell transfer as personalized immunotherapy for human cancer. Science 348, 62 LP – 68 (2015).

98. Sukumar, M., Kishton, R. J. & Restifo, N. P. Metabolic reprograming of anti-tumor immunity. Curr. Opin. Immunol. 46, 14–22 (2017).

99. Simoni, Y. et al. Bystander CD8+ T cells are abundant and phenotypically distinct in human tumour infiltrates. Nature 557, 575–579 (2018).

100. Scott, A. C. et al. TOX is a critical regulator of tumour-specific T cell differentiation. Nature 571, 270–274 (2019).

101. Khan, O. et al. TOX transcriptionally and epigenetically programs CD8+ T cell exhaustion. Nature 571, 211–218 (2019).

102. Alfei, F. et al. TOX reinforces the phenotype and longevity of exhausted T cells in chronic viral infection. Nature 571, 265–269 (2019).

103. Yao, C. et al. Single-cell RNA-seq reveals TOX as a key regulator of CD8+ T cell persistence in chronic infection. Nat. Immunol. 20, 890–901 (2019).

104. Seo, H. et al. TOX and TOX2 transcription factors cooperate with NR4A transcription factors to impose CD8^+^ T cell exhaustion. Proc. Natl. Acad. Sci. 116, 12410 LP – 12415 (2019).

105. Liu, X. et al. Genome-wide analysis identifies NR4A1 as a key mediator of T cell dysfunction. Nature 567, 525–529 (2019).

106. Chen, J. et al. NR4A transcription factors limit CAR T cell function in solid tumours. Nature 567, 530–534 (2019).

107. Utzschneider, D. T. et al. T Cell Factor 1-Expressing Memory-like CD8(+) T Cells Sustain the Immune Response to Chronic Viral Infections. Immunity 45, 415–427 (2016).

108. Hartmann, F. J. et al. Scalable Conjugation and Characterization of Immunoglobulins with Stable Mass Isotope Reporters for Single-Cell Mass Cytometry Analysis. Methods Mol. Biol. 1989, 55–81 (2019).

109. Zunder, E. R. et al. Palladium-based mass tag cell barcoding with a doublet-filtering scheme and single-cell deconvolution algorithm. Nat. Protoc. 10, 316–333 (2015).

110. Hartmann, F. J., Simonds, E. F. & Bendall, S. C. A Universal Live Cell Barcoding-Platform for Multiplexed Human Single Cell Analysis. Sci. Rep. 8, 10770 (2018).

111. Hartmann, F. J. et al. Comprehensive Immune Monitoring of Clinical Trials to Advance Human Immunotherapy. Cell Rep. 28, 819–831.e4 (2019).

112. van der Windt, G. J. W., Chang, C.-H. & Pearce, E. L. Measuring Bioenergetics in T Cells Using a Seahorse Extracellular Flux Analyzer. Curr. Protoc. Immunol. 113, 3.16B.1–3.16B.14 (2016).

113. R Development Core Team. R: A Language and Environment for Statistical Computing. (2008).

114. Kotecha, N., Krutzik, P. O. & Irish, J. M. Web-Based Analysis and Publication of Flow Cytometry Experiments. Curr. Protoc. Cytom. 53, 10.17.1–10.17.24 (2010).

115. Behbehani, G. K., Bendall, S. C., Clutter, M. R., Fantl, W. J. & Nolan, G. P. Single-cell mass cytometry adapted to measurements of the cell cycle. Cytom. Part A 81 A, 552–566 (2012).

116. Grömping, U. Relative Importance for Linear Regression in R: The Package relaimpo. J. Stat. Softw. 17, 1–27 (2006).

117. Bannon, D. et al. Dynamic allocation of computational resources for deep learning-enabled cellular image analysis with Kubernetes. bioRxiv 505032 (2019) doi:10.1101/505032.

118. Wickham, H. ggplot2: Elegent Graphics for Data Analysis. (Springer-Verlag, 2016).

