## Supplementary figures for "Multiplexed Single-cell Metabolic Profiles Organize the Spectrum of Cytotoxic Human T Cells"

Affiliations:

Supplementary figures:

Supplementary Fig. 1: Assay-specific validation of heavy-metal conjugated metabolic antibodies.

Supplementary Fig. 2: Robustness and reproducibility of the scMEP approach.

Supplementary Fig. 3: Metabolic profiles recapitulate dynamic changes in metabolic pathway activity.

Supplementary Fig. 4: Coordination of metabolic remodeling in human T cells.

Supplementary Fig. 5: scMEP recapitulates metabolic differences of human naïve and memory T cells

Supplementary Fig. 6: Identification of metabolic phenotypes of immune cell lineages across human tissues.

Supplementary Fig. 7: Immunohistochemistry validation of metabolic antibodies.

Supplementary Fig. 8: Imaging-based analysis of metabolic states in human colorectal carcinoma.

Supplementary tables:

Supplementary Table 1: Metabolic antibodies and panels

Supplementary Table 2: Donor characteristics

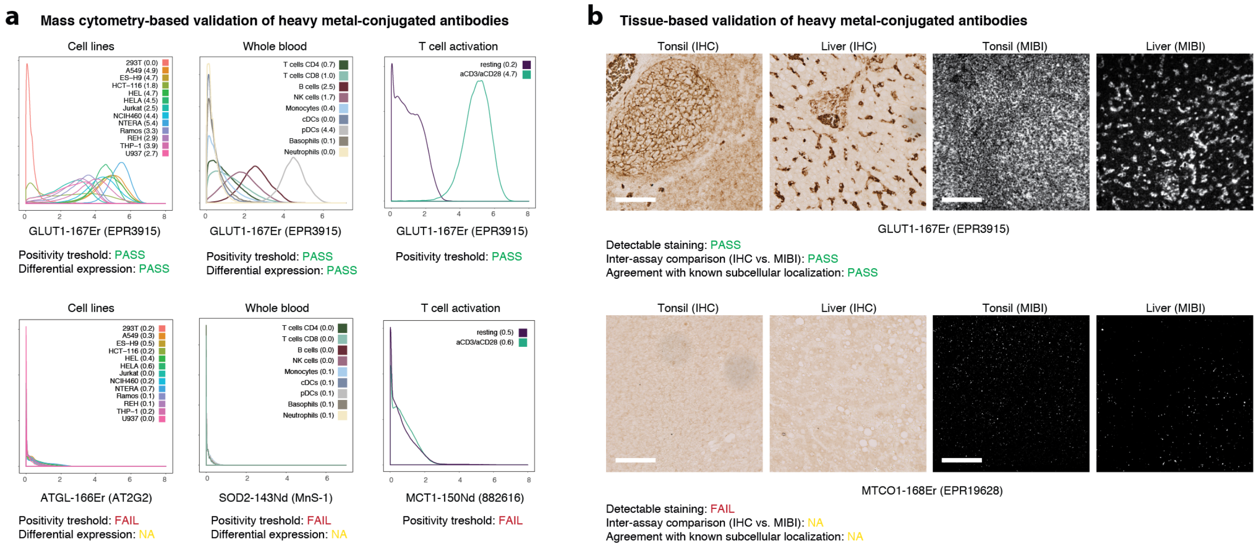


**Supplementary Fig. 1: Assay-specific validation of heavy-metal conjugated metabolic antibodies. a**, A broad range of cell lines was stained with heavy-metal conjugated antibodies and analyzed by mass cytometry (left). Human peripheral blood stained with lineage markers (CD45, CD3, CD4, CD8, CD45RA, CD66, CD14, CD19, CD20, HLA-DR, CD56, CD57, CD11c, CD123, FceRI, CD235ab) and metabolic antibodies to be validated (middle). Major immune cell types were identified using cell lineage markers and manual gating. Human T cells (resting and activated with anti-CD3/anti-CD38-beads for 72h) stained with metabolic antibodies (right). Numbers in brackets represent median arsinh values for the indicated population. Positive staining was defined as median > 10 ion counts, equal to asinh transformed value >1.5 of any subpopulation. Example of an antibody passing indicated quality control metrics (top) and range of examples of antibodies failing indicated metrics at various stages (bottom). Where available, cell-lineage specific expression and induction upon activation were compared to previously determined values (Geiger et al., 2016; Howden et al., 2019; Uhlen et al., 2015). **b**, Staining of control tonsil and liver FFPE tissues with metabolic antibodies analyzed through traditional IHC (left) and MIBI (right). Detectable staining was determined through visual inspection of both IHC and grayscale MIBI images. For intra-assay quality control, IHC and MIBI images were visually compared and in addition, related to previously determined staining patterns (Uhlen et al., 2015). Shown are examples of an antibody passing (top) or failing (bottom) the indicated quality control metrics. Scale bars = 100 μm. Results of this validation procedure for all antibodies are summarized in Supplementary Table 2.

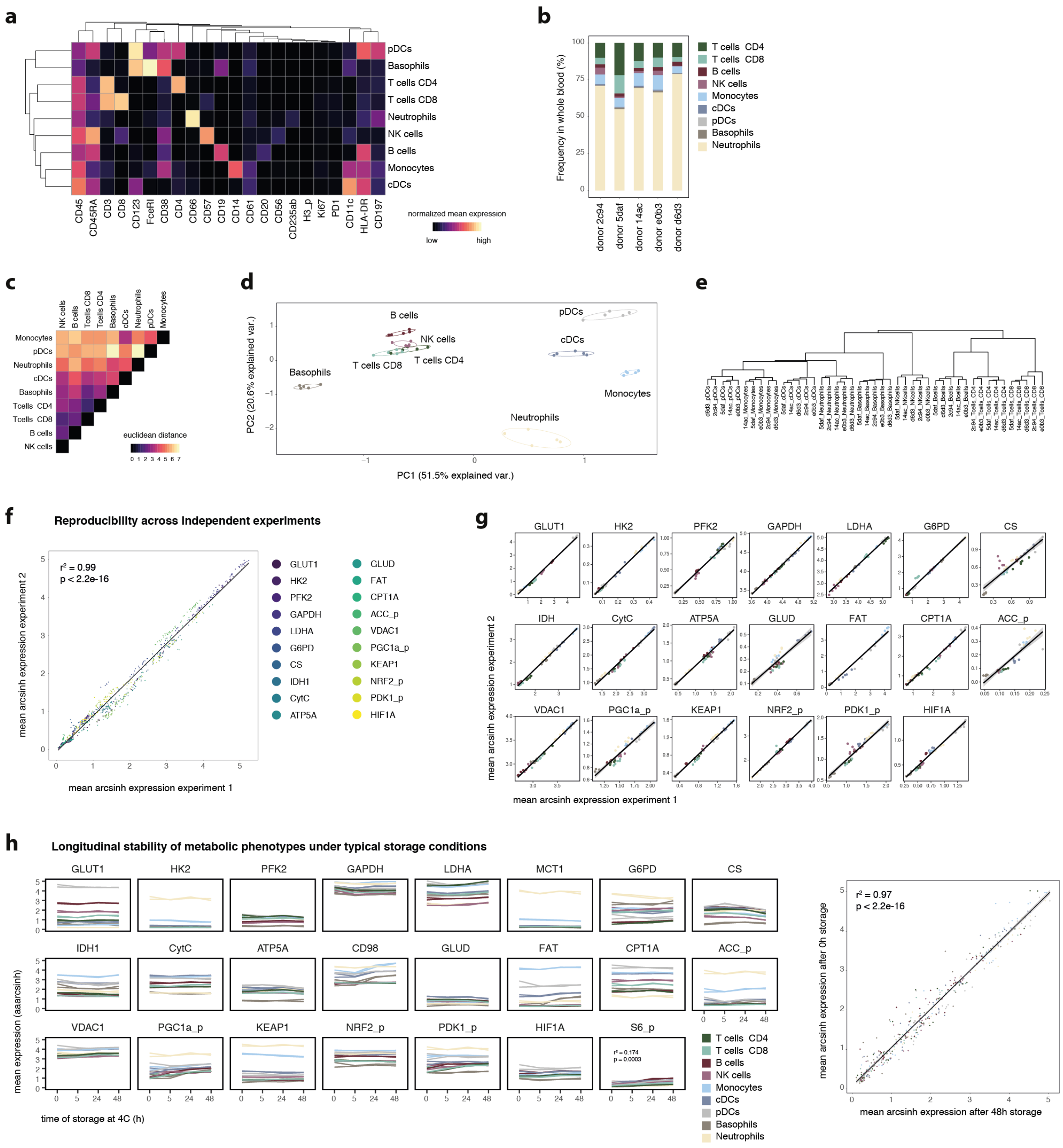

**Supplementary Fig. 2: Robustness and reproducibility of the scMEP approach.** Whole blood of healthy individuals (N = 5, see Supplementary Table 2) was fixed and stained with 23 metabolic and 22 immunological probes. **a**, Cell populations were identified through FlowSOM clustering and annotated into the major immune cell lineages. Shown here are mean normalized (99.9$^{th}$ percentile) expression values (arcsinh transformed) across identified peripheral immune cell populations. **b**, Frequencies of immune cells as identified in a across the five healthy donors. **c**, Euclidean distances between immune cell lineages based on the high-dimensional metabolic space (23 probes). **d**, First two components of a principal component analysis (PCA) of immune cell lineages as defined before, using only metabolic profiles and no immune markers. Each point represents one immune cell population from one healthy donor. Axes are scaled to the percentage of variance explained by the respective principal component. **e**, Hierarchical clustering (only using metabolic targets) of immune cell lineages from the five healthy individuals. **f**, Cells from the five healthy individuals as in a were stained and analyzed in two independent experiments using a highly similar panel and immune cell lineages were assigned as in a. Each point represents a mean arcsinh transformed value of one metabolic marker in one immune cell lineage. Points are colored by the respective metabolic probe. The black line as well as the $r^2$ and p-value show the results of a linear regression model. **g**, Mean arsinh values as in f for each marker separately. Colors indicate the respective immune cell population. **h**, Whole blood (two technical replicates) was drawn into a sodium heparin tube and either processed (red blood cell lysis and fixation) immediately or stored at 4 °C for up to 48 h before processing. All samples were then barcoded into a composite sample and analyzed as before. Charts show mean arcsinh-transformed values for each metabolic marker on a given immune cell lineage across time (left). Linear regression between mean population values of the 0 and 48 h storage times. The black line as well as the $r^2$ and p-value show the results of a linear regression model.

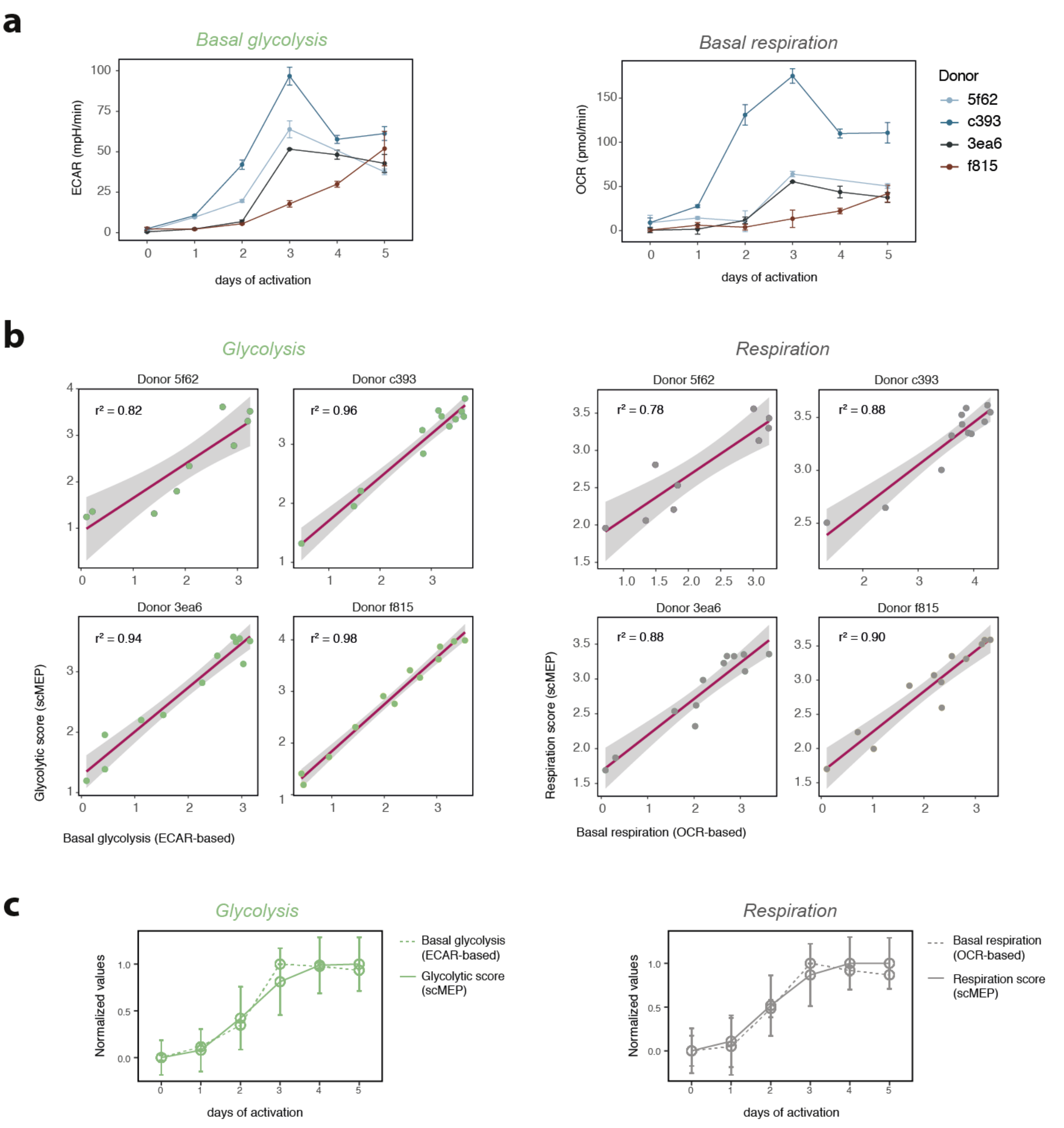


**Supplementary Fig. 3: Metabolic profiles recapitulate dynamic changes in metabolic pathway activity. a**, PBMCs were isolated from healthy donors (N = 4). Naïve or memory T cells (containing $CD4^{+}$ and $CD8^{+}$ cells) were purified by negative isolation with magnetic beads. Purified T cell populations (naïve or memory) were then activated using anti-CD3/anti-CD28 beads for 0-5 days. Shown are extracellular flux analysis-derived basal glycolysis (ECAR-based, left) and basal respiration (OCR-based, right) across different days of activation and across multiple independent donors. **b**, Mass cytometry-based scMEP scores (glycolysis left, respiration right) represent the mean (scaled) expression of all metabolic targets within a given pathway. Each dot represents the mean scMEP score of a T cell population (naïve or memory) stimulated with anti-CD3/anti-CD28 for 0-5 days. Both values were arcsinh transformed (cofactor 5) for this analysis. Black lines and $r^2$ values represent results of a linear regression model, with black shading representing the 95% CI. **c**, Normalized (minimum to maximum scaled) glycolytic (left) and respiration (right) scores across days of stimulation. Error bars represent normalized s.d. in between technical replicates for extracellular flux analysis-based values and between single-cells for scMEP values.

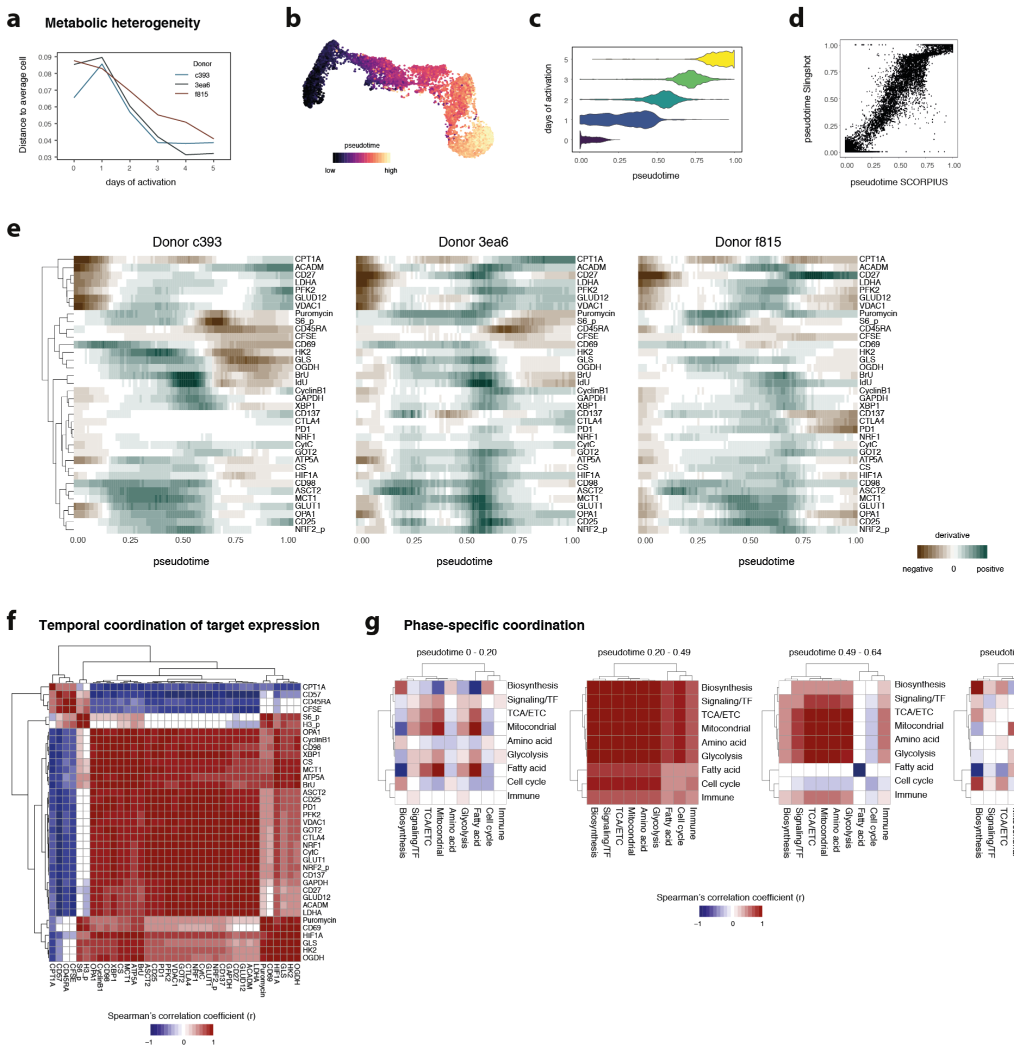


**Supplementary Fig. 4: Coordination of metabolic remodeling in human T cells.** PBMCs were isolated from healthy individuals (N = 3). Naïve T cells (containing $CD4^{+}$ and $CD8^{+}$ cells) were purified by negative isolation with magnetic beads. Purified T cell populations were then activated using anti-CD3/anti-CD28 beads for 0-5 days. **a**, Mean metabolic heterogeneity within naïve human $CD8^{+}$ T cells was calculated as cosine distance (based on all metabolic markers except IdU, but no phenotyping markers) to an average cell within the given day of activation. **b**, Cells on a two-dimensional UMAP projection of the high-dimensional space were colored according to their SCORPIUS-inferred pseudotime. **c**, Distribution of cells from different days of activation across pseudotime. **d**, The same cells and the same markers were used to infer pseudotime using an independent algorithm (Slingshot). **e**, Data was binned into 100 bins and averaged for each bin. Slope (first derivative) of marker expression across pseudotime for three independent donors analyzed in three independent experiments. **f**, Binned data as in e was used as an input for Spearman's correlation analysis. P-values were BH-adjusted to correct for multiple hypothesis testing and r values were set to 0 for all BH-corrected P values > 0.05. **g**, Spearman's r as in f stratified by pseudotime bins based on the previously identified inflection points.

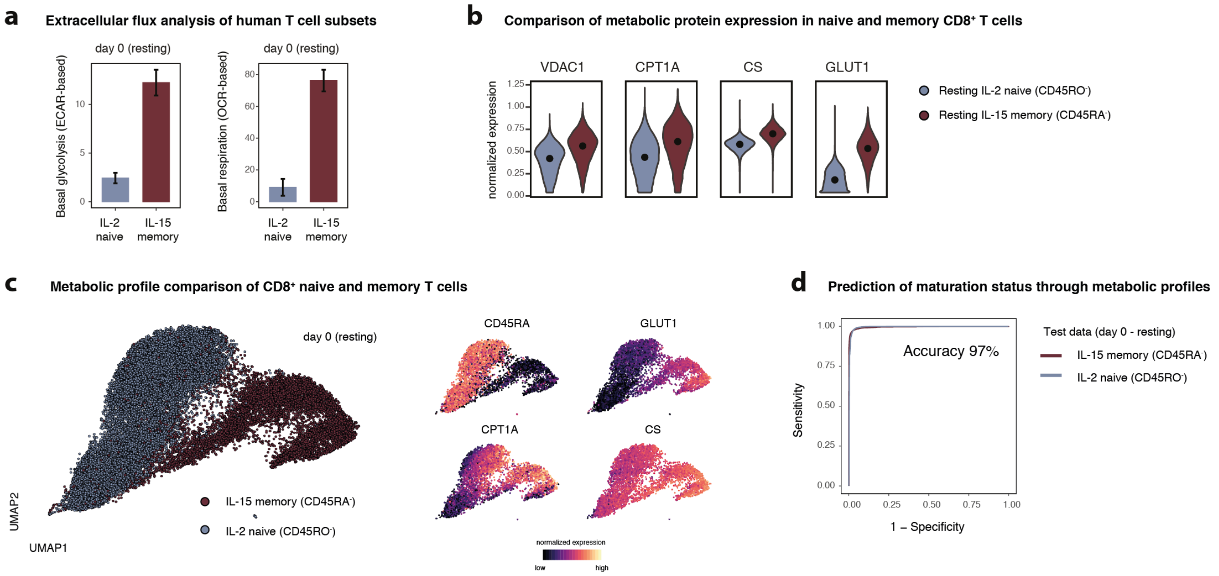


**Supplementary Fig. 5: scMEP recapitulates metabolic differences of human naïve and memory T cells.** PBMCs were isolated from healthy individuals (N = 3). Naïve T cells (containing CD4$^{+}$ and CD8$^{+}$ cells) were purified by negative isolation with magnetic beads. Purified T cell populations were then activated using anti-CD3/anti-CD28 beads for 0-5 days. **a**, Extracellular flux analysis of resting (day 0) IL-2 naïve (CD45RO$^{-}$) and IL-15 memory (CD45RA$^{-}$) T cells (containing CD4$^{+}$ and CD8$^{+}$). ECAR-based basal glycolysis (left) and OCR-based basal respiration (right). Shown are cells from one representative donor. **b**, Examples (one representative donor) of metabolic differences between naïve and memory CD8$^{+}$ T cells as in a. **c**, Cells were subsampled for equal representation of naïve and memory CD8$^{+}$ T cells. Two-dimensional UMAP projection of the metabolic space with cells colored by their maturation status (left) or colored by their normalized expression level of markers as in g (right). **d**, L1 regularized linear regression (using only metabolic profiles) was trained on a subset of cells (resting, 0 days of activation) from one donor and tested on a separate set of cells from the same donor. Shown are results from the test dataset and the overall accuracy of the model.

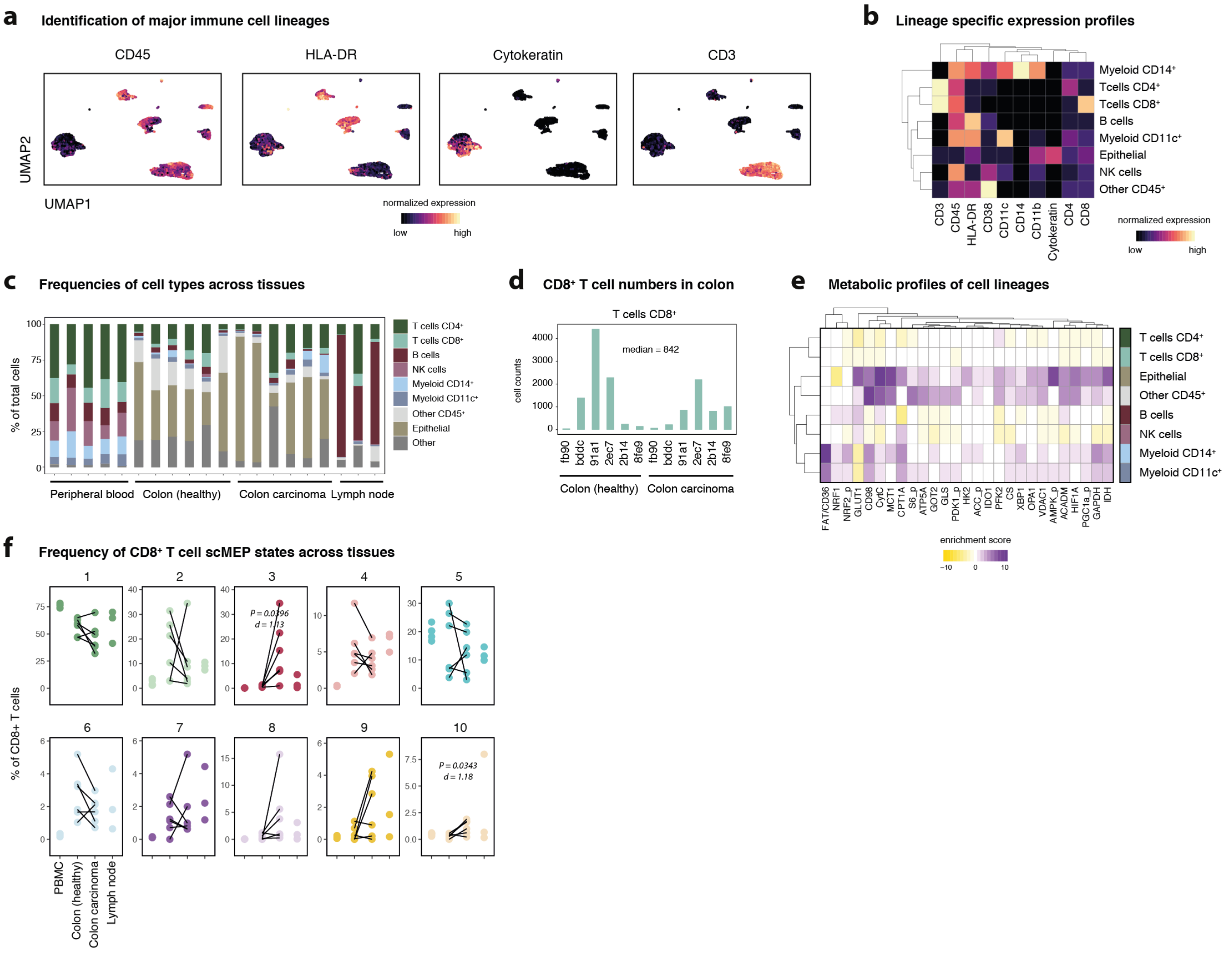


**Supplementary Fig. 6: Identification of metabolic phenotypes of immune cell lineages across human tissues.** Healthy donor PBMC (N = 5), lymph node biopsies (N = 3) as well as single-cell suspensions from colorectal carcinoma (N = 6) and matched adjacent healthy sections (N = 6, see Supplementary Table 2) were barcoded, stained and acquired on a mass cytometer. **a**, Major cell lineages from all samples and tissues were identified through FlowSOM-based clustering. UMAP-dimensionality reduction was calculated using subsampled data from all lineages and all available markers. Cells are colored by their normalized expression value of the indicated lineage markers. **b**, Mean normalized expression values of the major lineage markers across cell populations. **c**, Frequencies of cell populations across all samples. **d**, Total counts of $CD8^+$ T cells from colon samples. **e**, Marker enrichment modeling (MEM) of metabolic phenotypes across the major cell lineages. **f**, Statistical analysis of frequencies of scMEP phenotypes within $CD8^+$ T cells. P-value was calculated using a pairwise t-test between healthy and malignant sections from the same patient. Effect size is represented as Cohen's d. P-values not stated when P > 0.05.

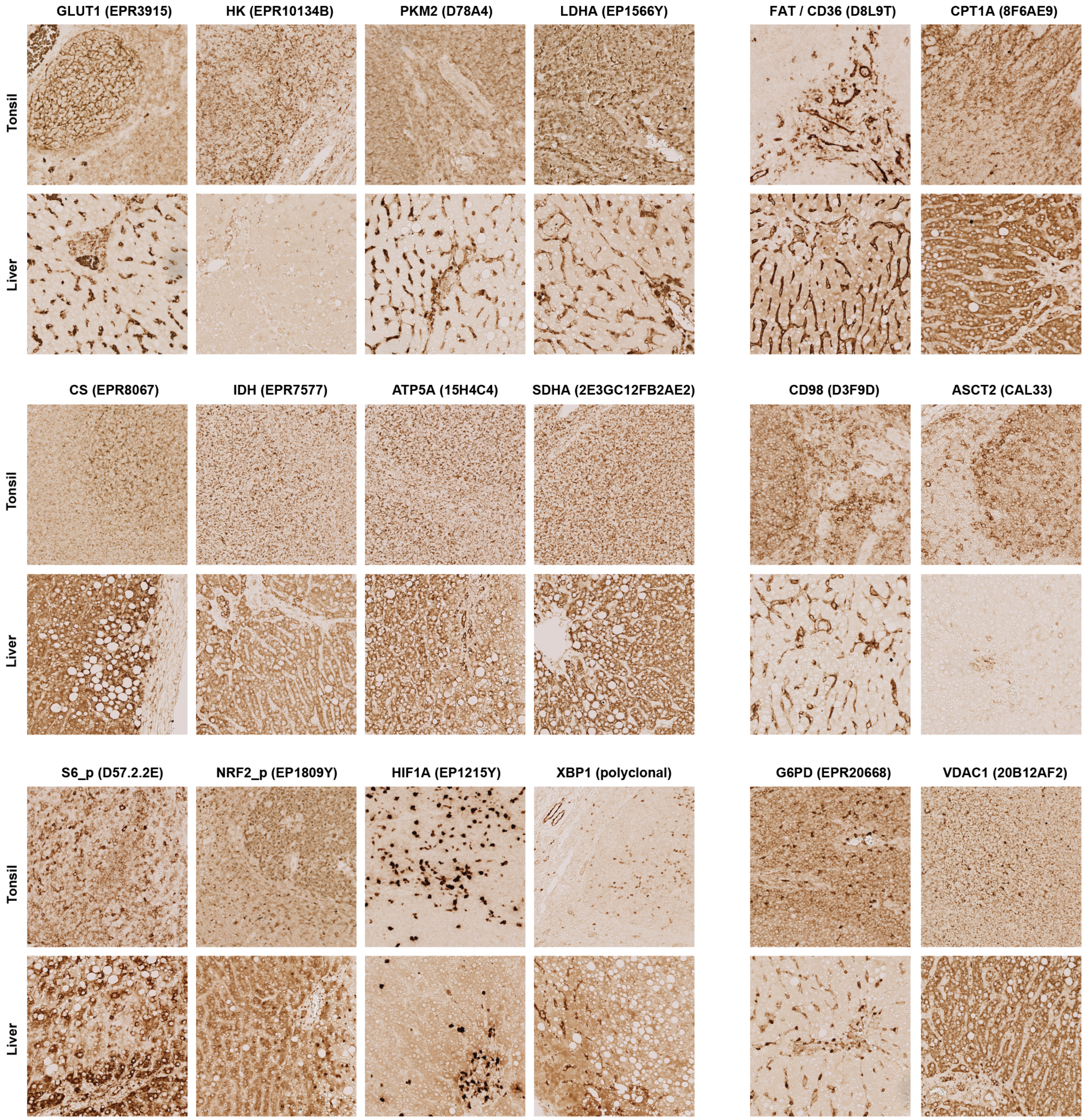


**Supplementary Fig. 7: Immunohistochemistry validation of metabolic antibodies.** FFPE sections from healthy donor liver and tonsil tissues were stained by IHC with the indicated (metal-conjugated) antibodies for validation.

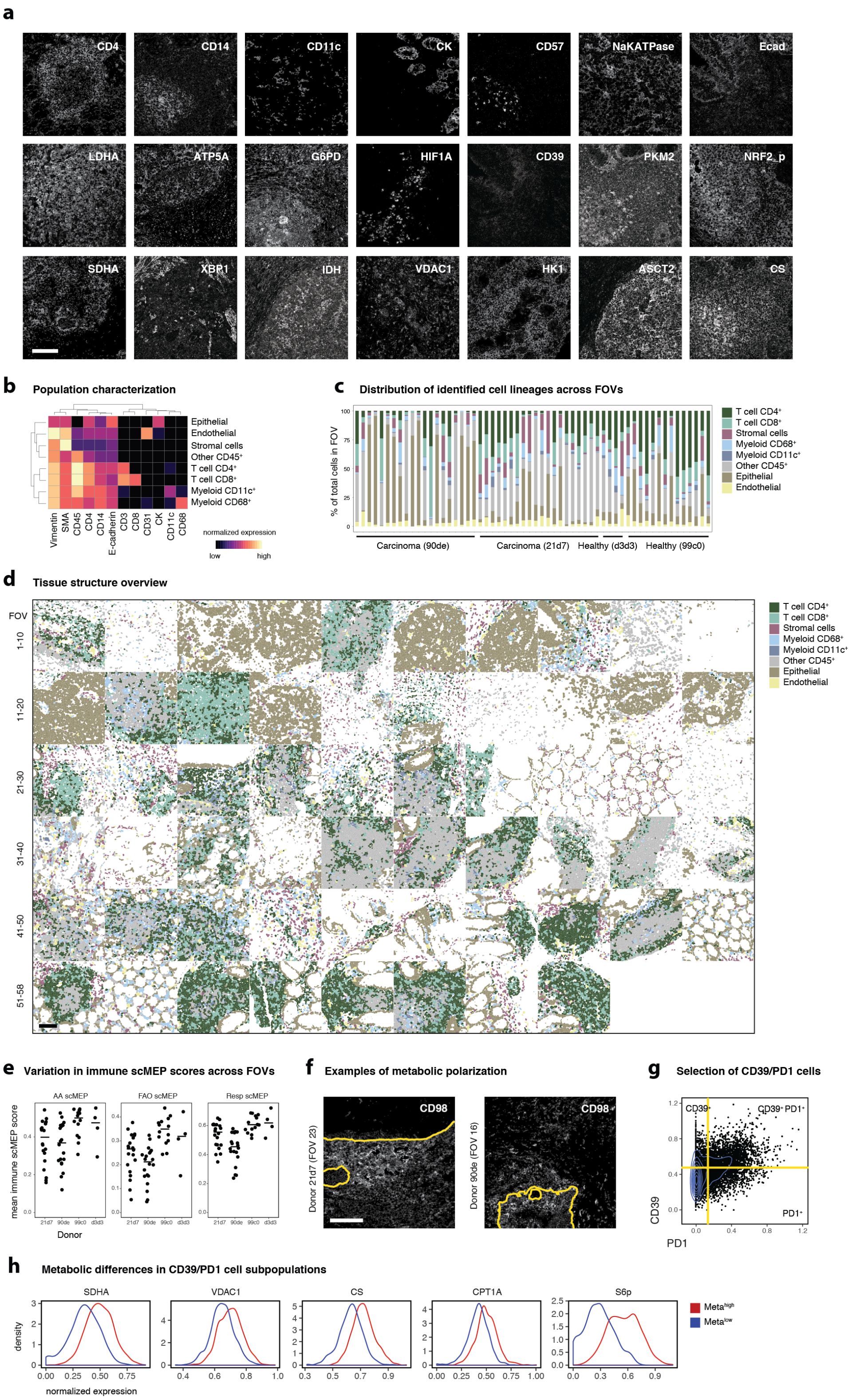


**Supplementary Fig. 8: Imaging-based analysis of metabolic states in human colorectal carcinoma. a**, Exemplary grayscale images showing staining of lineage and metabolic antibodies in human colorectal FFPE sections imaged by MIBI. Scale bar = 100

μm. **b**, Single cells were segmented from all images and clustered into the main cell lineages using FlowSOM. Heatmap values represent scaled and normalized mean expression values of the indicated population. **c**, Composition of each field of view (FOV) based on clustering results as in b. FOVs are numbered 1-58 from left to right and top to bottom. **d**, Segmented and clustered cells can be visualized in their original location on the image. **e**, Average scMEP scores for all $CD45^+$ cells within a FOV. AA = Amino acid. FAO = Fatty acid oxidation. Resp = respiration. **f**, Exemplary grayscale images of two FOVs showing polarization of CD98 towards the tumor immune border, indicated in yellow. Scale bar = 100 μm. **g**, CD39 and PD1 expression on $CD8^+$ T cells across all FOVs. CD39/PD1 cells were defined using the indicated yellow lines. **h**, CD39/PD1 cells defined as in g were clustered into two subsets using FlowSOM and their metabolic markers as input. Histograms display the marker expression levels of the two clusters, termend $Meta^{high}$ (red) and $Meta^{low}$ (blue).
